# Translation Accuracy in *E*. *coli*

**DOI:** 10.1101/2025.04.18.649569

**Authors:** Ryan Stikeleather, Farhan Ali, Wei-Chin Ho, Tim Licknack, Michael Lynch

## Abstract

We present a method that quantifies nearly all pairwise amino acid substitutions, revealing the rates and spectra of mistranslation per amino acid and per codon. While determining error rates is illuminating, understanding why certain mistranslation events are enriched requires further exploration, particularly in relation to the translation-accuracy hypothesis. We found that codons favored in highly expressed genes are not predictive of translational fidelity and are not selected for translational accuracy. We evaluated our method using historically characterized ribosomal variants of *Escherichia coli* reported to differ in translation fidelity. However, we found no significant differences among their overall translation-error rates, estimated here to be ∼2 per 1,000 amino acids. Instead they each exhibited unique mistranslation profiles, which likely led to prior overestimates of the translation error-rate in the putative error-prone variant. Together, these results underscore the significance of a direct, proteome-wide measure of translation fidelity.

## Introduction

The total translation-error rate of an organism is the composite of the rates associated with ribosomal misreading and tRNA mischarging. Estimates of fidelities for both misreading and mischarging are similar, on the order of 10^-4-10-3^ per codon, which is at least an order of magnitude higher than the transcript-error rate^1–4^. In some cases, mischarging rates have even been reported to be on the order of 10^-2 5,6^. However, methods for measuring translation-error rates have primarily relied on a single codon, or even single nucleotide challenges as in the case of dipeptide formation experiments and reporter assays^2,7^. These approaches fail to capture the entire spectrum of potential amino-acid misincorporations. A need for improved measures of the translation-error rate was expressed fifteen years ago, and yet, as of this writing, only two other groups have attempted to use mass spectrometry to determine the entire spectrum of translation errors^8–10^.

In this work, we present total translation-error rates for three *E*. *coli* strains (a generous gift from Dr. Hani Zaher), each carrying a distinct ribosomal mutation that has been previously characterized to affect fidelity of translation^2,11^. These *E*. *coli* strains are all derived from a Xac *E*. *coli* (*ara,* Δ*lacproAB gyrA, rpoB, argE*[amber]), otherwise considered here to be the wild-type variant, with an unmodified ribosome. The restrictive (*res*) ribosomal variant, with putatively improved accuracy, contains a K43N mutation in RpsL, an amino-acid site that contacts a decoding nucleotide C912^2^. The putative error-prone ribosomal variant, or the ribosomal ambiguity (*ram*) strain, contains a 5 nt-deletion in *rpsD* from nucleotides 528-532, resulting in a 180 amino-acid truncation of the S4 protein that usually interacts with S5^2^. Originally, the accurate *E*. *coli* was isolated among many others as a streptomycin-resistant mutant. Interestingly, the most resistant of these isolated revertants also became dependent on streptomycin for growth, and the error-prone ribosomal variant was isolated as a revertant of streptomycin dependence^11–16^.

Both dipeptide formation and dual-reporter luciferase assays were employed to originally characterize these *E*. *coli* strains as being accurate or error-prone^2,11,17^. Dipeptide formation is carried out *in vitro* with short synthetic mRNA, usually less than 60 nt in length and consisting of only two or three different amino acids. The specific sequence characteristics of the mRNA and the specific *in vitro* buffer system utilized during the experiment might affect error estimates^18^. Moreover, like binary competition experiments, dipeptide formation assays only employ two presumably pure and correctly charged tRNAs, whereas *in vivo* all tRNAs for the twenty amino acids are competing for the ribosome^5,6^. Reporter assays capture more factors important to translation, such as the previously mentioned *in vivo* competition of tRNAs, but they are still limited to the evaluation of one codon of one protein. In the case of the luciferase dual-reporter system, evaluation of the translation error is dependent on the misreading of an Asn AAU codon under the presumption that a misread would most likely incorporate a Lys (AAA). However, in firefly luciferase, 21 amino-acid sites affect the luminescence of the protein either positively or negatively dependent on the substitution^19^. Of course, any errors that do not improve luminescence will be undetected.

In addition to these technical limitations, the usage of reporter assays is also restricted to species that can be genetically manipulated. Here, we present a straightforward method that should be widely applicable for determining translation-error rates, the spectrum of amino-acid and codon substitution rates, and lastly protein- and codon-level error rates.

## Results

### Amino-acid substitution spectra reveal bias in mistranslation events

The method described here employs a simple LC-MS workflow and a database search to identify wild-type and mistranslated peptides. Purified proteins are subjected to peptide preparation, which includes treatment with trypsin that results in predictable peptide fragments. An initial search using the wild-type proteome is first carried out to identify proteins that are present in the sample. This information is used to generate a custom database of wild-type peptides and potentially mistranslated peptides. This step involves performing an *in silico* trypsinization of the wild-type protein sequences and subsequent *in silico* mutagenesis, wherein all possible singly substituted amino-acid variants are recorded. The custom database is then used as a platform for interrogating the mass spectrometry data. Peptide spectral matches are validated using a target false-discovery rate (FDR) and a posterior error probability (PEP) cutoff of 1%^20,21^. Validated peptide spectral matches are processed through a custom bioinformatics pipeline to generate the rate and spectra of amino-acid substitutions (Figure 1A).

**Figure 1.**
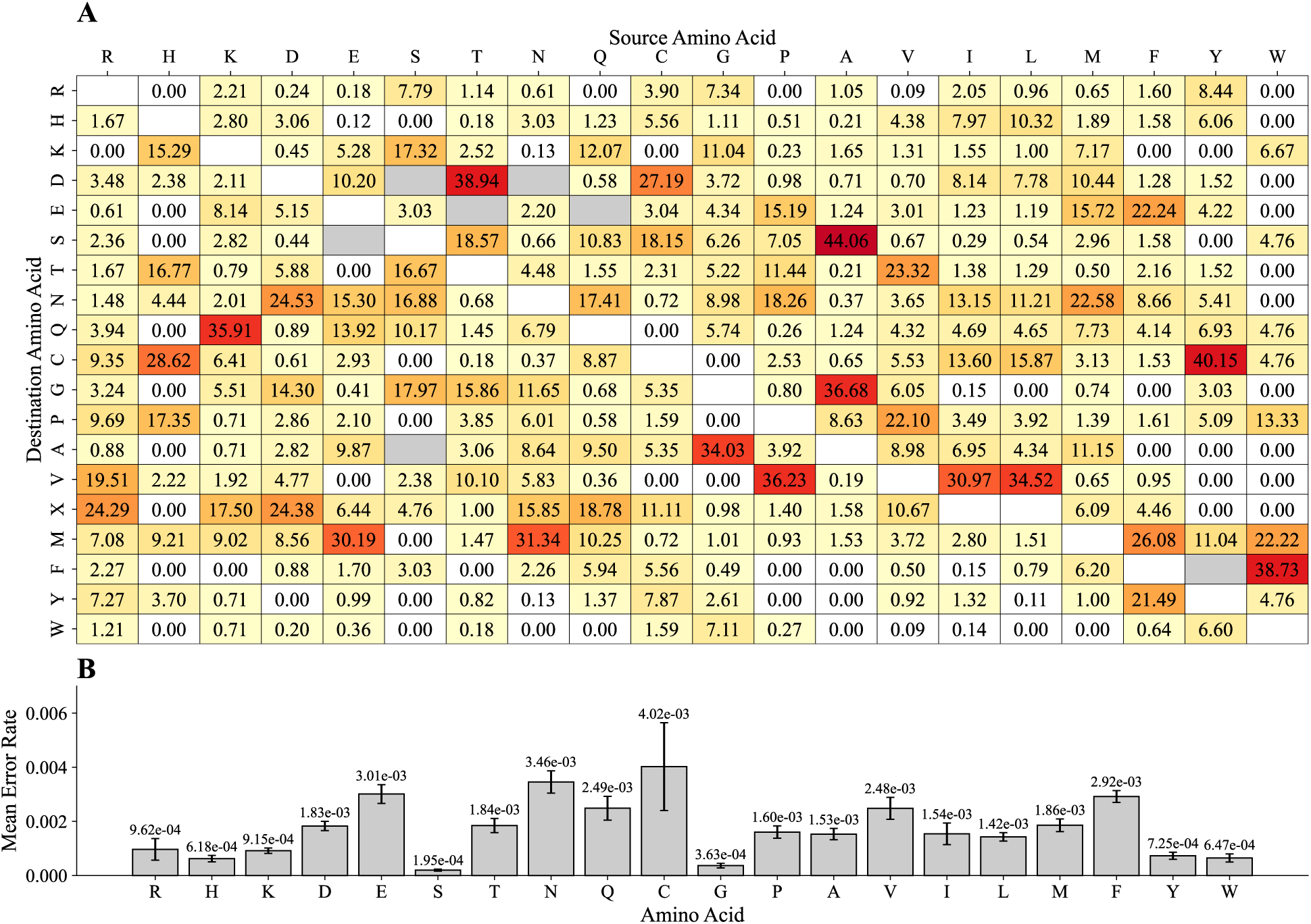
Mean rate and spectrum of translation errors of wild-type *E*. *coli*. A) Wild-type (source) amino acids are represented as columns, and erroneously incorporated (destination) amino acids are represented as rows. B) The mean error rate at which each amino acid is mistranslated is plotted with standard error bars across three biological replicates. Substitutions that are indistinguishable from chemical modifications of the source amino acid (see Table 3) are removed from the analysis (grey boxes). Leucine and isoleucine are combined in the destination row “X.” Subfigure B gives the average rate at which each amino acid is mistranslated, while subfigure A gives the percentage distribution of those mistranslations across destination amino acids.

Wild-type *E*. *coli* exhibited a mean translation error rate of 1.82 × 10^-3^ per codon (standard error = 5.92 × 10^-5^). The mean error rate for each amino acid, defined as the number of detected substitutions divided by the total number of sites sampled, is reported in Figure 1B. The rates of mistranslation vary considerably across amino acids (χ2 = 1585.63, *P* < 0.001). Notably, serine and glycine exhibited significantly lower error rates compared to other amino acids, while glutamic acid and asparagine experienced significantly higher error rates. The molecular details in the collected data also allow us to quantify the error rates per codon (Figures S3-S5) or per protein (Figure 4A).

To further study the spectra of translation errors in wild-type *E*. *coli*, we performed a statistical test on the observed number of substitutions using simulated distributions of expected substitutions based on the marginal probabilities of each source and destination amino acid (Methods). In an attempt to explain why these substitutions were occurring more frequently than would be expected by chance, the list of the top 5% of substitutions is summarized in Table 1 (Complete data in Table S1). The minimum mutational distance (MMD) is the minimum number of codon positions that would need to be misread to observe any particular amino acid substitution. For example, methionine (ATG) to arginine (AGG) mistranslation requires a single base to be misread at position 2 of the codon, resulting in an MMD of 1. An MMD of one nucleotide likely results from ribosomal decoding errors, while an MMD of 3 nucleotides suggests tRNA mischarging, as ribosomes are unlikely to misread three nucleotides. A two nucleotide MMD is likely some combination of both decoding and mischarging error.

**Table 1.** Mistranslation events in the top 5% of Z-scores for the wild-type ribosomal variant.

| Amino Acid Source | Amino Acid Destination | Z-score | Minimum Distance (nt) |
| --- | --- | --- | --- |
| A | S | 35.74 | 1 |
| F | Y | 26.29 | 1 |
| T | D | 24.47 | 2 |
| A | G | 21.33 | 1 |
| V | P | 20.23 | 2 |
| L | V | 19.63 | 1 |
| V | T | 16.50 | 2 |
| G | A | 16.11 | 1 |
| K | Q | 15.79 | 1 |
| E | M | 15.48 | 2 |
| F | E | 15.18 | 3 |
| I | V | 14.82 | 1 |
| L | C | 14.20 | 2 |
| W | F | 13.72 | 2 |
| N | M | 13.65 | 2 |
| P | V | 13.62 | 2 |
| D | X | 11.44 | 2 |
| G | W | 11.21 | 1 |

The significance (nominal $P$ -value $\leq 0.001$ and top 5% Z-score of observation) was determined by comparing the observed number of substitutions to the distribution of 10,000 simulated values of expected substitutions, dependent on the marginal probabilities of sources and destinations. The distance was defined by the minimum number of nucleotides per codon required to be misread to cause the observed translation error.

Using similar methods, we also quantified the rates and spectra of translation errors for the “accurate” and “error-prone” ribosomal variants, respectively (Figures S1 and S2). The mean translation-error rate for the wild-type (1.82 × 10^-3^, SE = 5.92 × 10^-5^) was similar to that of the “accurate” ribosomal variant (1.86 × 10^-3^, SE = 1.83 × 10^-4^). In comparison, the “error-prone” ribosomal variant exhibited a mean error rate approximately 20% higher (2.23 × 10^-3^, SE = 2.19 × 10^-4^). However, the differences in overall translation-error rates among the three variants were not statistically significant (one-way ANOVA, *P* = 0.17).

### The “error-prone” ribosomal variant misreads the third position of codons, with an Asn to Lys bias

Decoding errors can be caused by ribosomal misreading at any of the three nucleotide positions within the codon. Inferring the likely position of each error, based on the nature of the genetic code, provides insight into position-specific decoding fidelity. The data show that the error-prone ribosomal variant misreads codons at the third position significantly more often than the wild-type or accurate ribosomal variants (*P* = 0.0009, one-way ANOVA) (Figure 2A). Additionally, when missense substitutions occur, the MMD of the error-prone ribosomal variant is significantly more likely than the wild-type or accurate ribosomal variants to be only one nucleotide (*P* = 0.01, one-way ANOVA) (Figure 2B).

**Figure 2.**
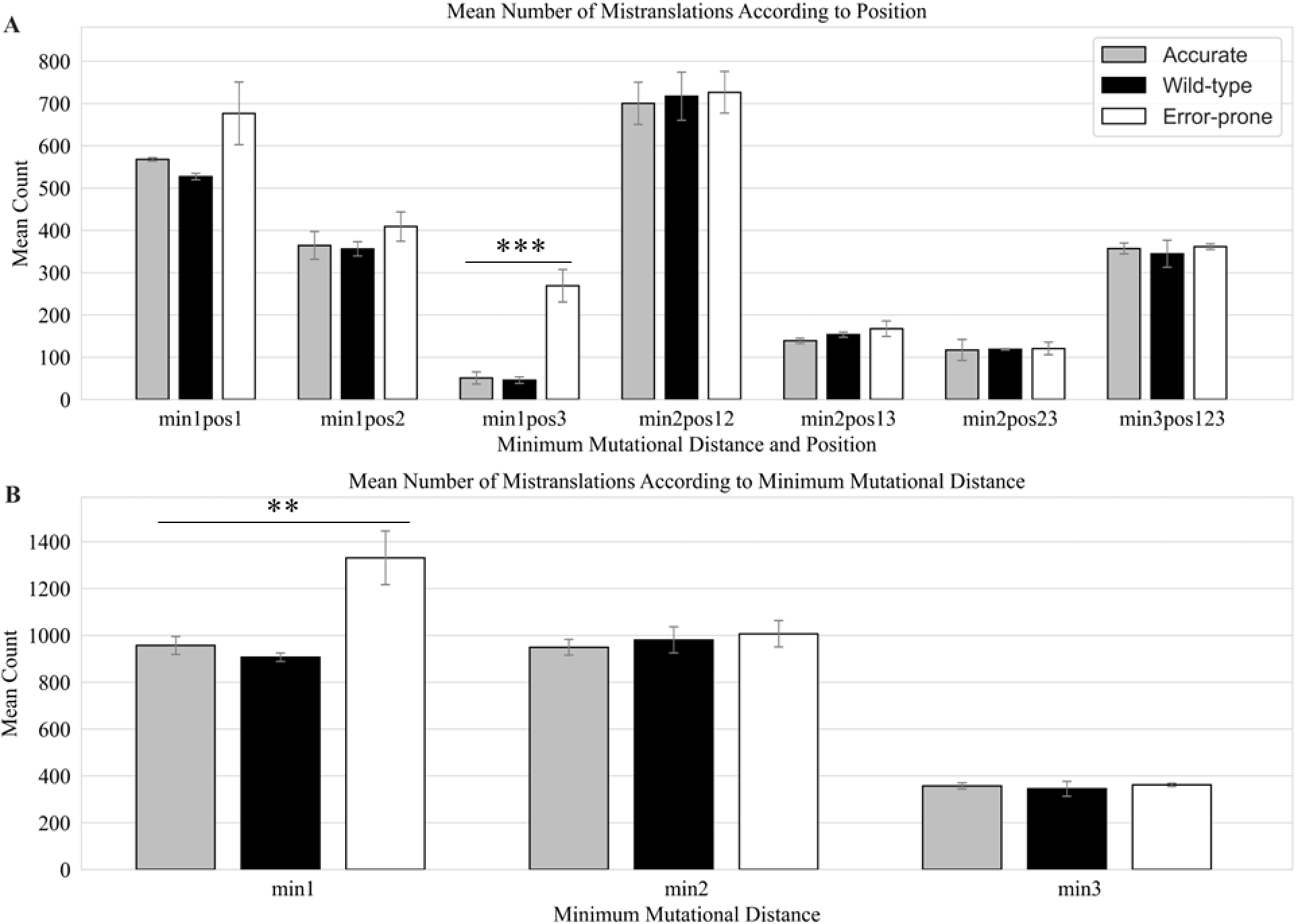
Mistranslation events according to minimum mutational distance (MMD) and position. A) Each ribosomal variant is color-coded according to the legend. The categories start with “min” followed by a number, indicating the MMD. For each MMD, the position is indicated by “pos” followed by the number(s). Codon position is left to right, 1 to 3. Error bars represent the standard error of the mean. Asterisks are indicative of statistically significant differences, \*\**P* ≤0.01 and \*\*\**P* ≤0.001. B) Mean number of mistranslation events according to MMD.

Historically, AAU asparagine has been one of the most studied codons in relation to translation-error rates, as it has commonly been used as a near-cognate challenge codon at the active site of luciferase. Translation-fidelity studies have used it because it differs by only a single nucleotide in comparison to AAA lysine, which is the wild-type codon at the active site of luciferase. In fact, the *E*. *coli* ribosomal variants used in this study were previously characterized via luciferase reporter assay^2,7^. When comparing the mean error rates of the AAU codon between the wild-type, accurate, and error-prone variants here, the rate for the error-prone variant was approximately double that of the other two ribosomal variants (5.62 × 10^-3^ vs. 2.99 × 10^-3^ and 3.30 × 10^-3^). More importantly, 52% of all mistranslations at AAU codons in the error-prone ribosomal variant, on average, were to lysine (Figure S5), whereas for the wild-type and accurate variants lysine was never erroneously incorporated at AAU codons (Figures S3, S4). Thus, the data clearly show that the modified ribosomes behave differently. However, the combination of a higher error rate at the AAU codon and a bias toward incorporating lysine—the amino acid enabling luminescence in the reporter assay—likely led to prior overestimates of the translation-error rate of the error-prone variant. When third-position AAU mistranslation events are removed from the analysis, the average number of errors in the third-position remains significantly higher than the wild-type and accurate ribosomal variants (*P* = 0.003, one-way ANOVA). Therefore, the “error-prone” variant mistranslates the third position significantly more often than the other two variants, a difference that cannot be attributed to AAU mistranslation alone.

### Codons favored in highly expressed genes are translationally inaccurate

Synonymous codons are used unequally across the *E*. *coli* genome. Frequencies of some codons increase with gene expression, and correlate with the cellular abundance of their cognate tRNAs, leading to the suggestion that these codons are favorable to the process of translation^22–24^. The standard model of “translational selection” considers that a set of codons are preferred for their translational speed as well as accuracy^25^. However, a direct test of this hypothesis has been lacking due to an absence of estimates for genome-wide translation error rates of all codons. Instead, codons thought to be preferred by translational selection in *E*. *coli* were initially identified based on their observed frequencies in a small subset of genes^26^. A concern here is that because the intrinsic mutational biases of a genome also shape its codon composition, the most frequent codon for an amino-acid might not necessarily be favored by selection^24,27^.

We investigated whether codons favored in highly-expressed genes are translated accurately based on our estimates of codon-level mistranslation rates. To identify preferred codons, we analyzed coding sequences of a set of 429 highly abundant proteins (top 25% proteins based on protein-abundance data from PaxDB)^28^. Unlike previous studies that assumed the most frequent synonymous codons were preferred codons, we identified selectively favored codons based on a quantitative model that takes mutational biases into account^29,30^. The model, detailed in Methods (Identification of Preferred Codons), compares observed frequencies of synonymous codons to their expected frequencies given mutation rates between synonymous codons. To correct for mutational bias, we used base-pair substitution rates derived from a mutation accumulation study of *E*. *coli* PFM2^31^. Using this approach, we estimated codon-specific selection coefficients for the 18 amino acids with more than one synonymous codon. For each amino acid, the codon with the highest selection coefficient among synonymous codons was considered as the preferred codon. The resulting set of preferred codons predicted (Table 2 – Column 1) matched the one suggested by Sharp and Li^26^ (Table 2 – Column 3), that were defined based on the observed codon frequencies of only 27 highly expressed genes, for all except three amino acids - Ala, Ser, and Gly.

**Table 2.** Selectively preferred codons as determined by the Li-Bulmer model of mutation-selection-drift equilibrium (Li 1987, Bulmer 1991) applied to codon frequencies in the top 25% of genes by protein abundance (PaxDb) or, alternatively, all genes.

| Amino acid | (1) Top 25% | (2) All | (3) SL87 |
| --- | --- | --- | --- |
| A | GCG | GCG | GCT |
| C | TGC | TGC | TGC |
| D | GAC | GAT | GAC |
| E | GAA | GAA | GAA |
| F | TTC | TTC | TTC |
| G | GGC | GGC | GGT |
| H | CAC | CAT | CAC |
| I | ATC | ATC | ATC |
| K | AAA | AAA | AAA |
| L | CTG | CTG | CTG |
| N | AAC | AAC | AAC |
| P | CCG | CCG | CCG |
| Q | CAG | CAG | CAG |
| R | CGT | CGC | CGT |
| S | AGC | AGC | TCT |
| T | ACC | ACC | ACC |
| V | GTT | GTG | GTT |
| Y | TAC | TAC | TAC |

The four rows in yellow are for amino acids where preferred codons differ between the two sets of genes. Preferred codons identified in Sharp & Li 1987 (SL87) are listed in the last column for comparison. The three amino acids where these differ from the top 25% set are in orange. Preferred codons remain the same across all columns for the remaining eleven amino acids in blue.

We found a strong positive correlation between the average frequency of preferred codons among synonymous codons in a gene and the abundance of its protein product (Figure 3A). Furthermore, the relative expression of cognate tRNAs, based on a published RNA-Seq dataset^32,33^, was greater for preferred codons compared to that of their synonyms (Figure S6). Upon considering all genes instead of only the highly expressed ones, we identified a slightly different set of preferred codons (Table 2 – Column 2). However, we did not see an enrichment of preferred codons with increasing protein abundance for this alternative set of preferred codons (Figure S7). Together, these observations suggest that the codons selectively favored in highly expressed genes are under translational selection.

**Figure 3.**
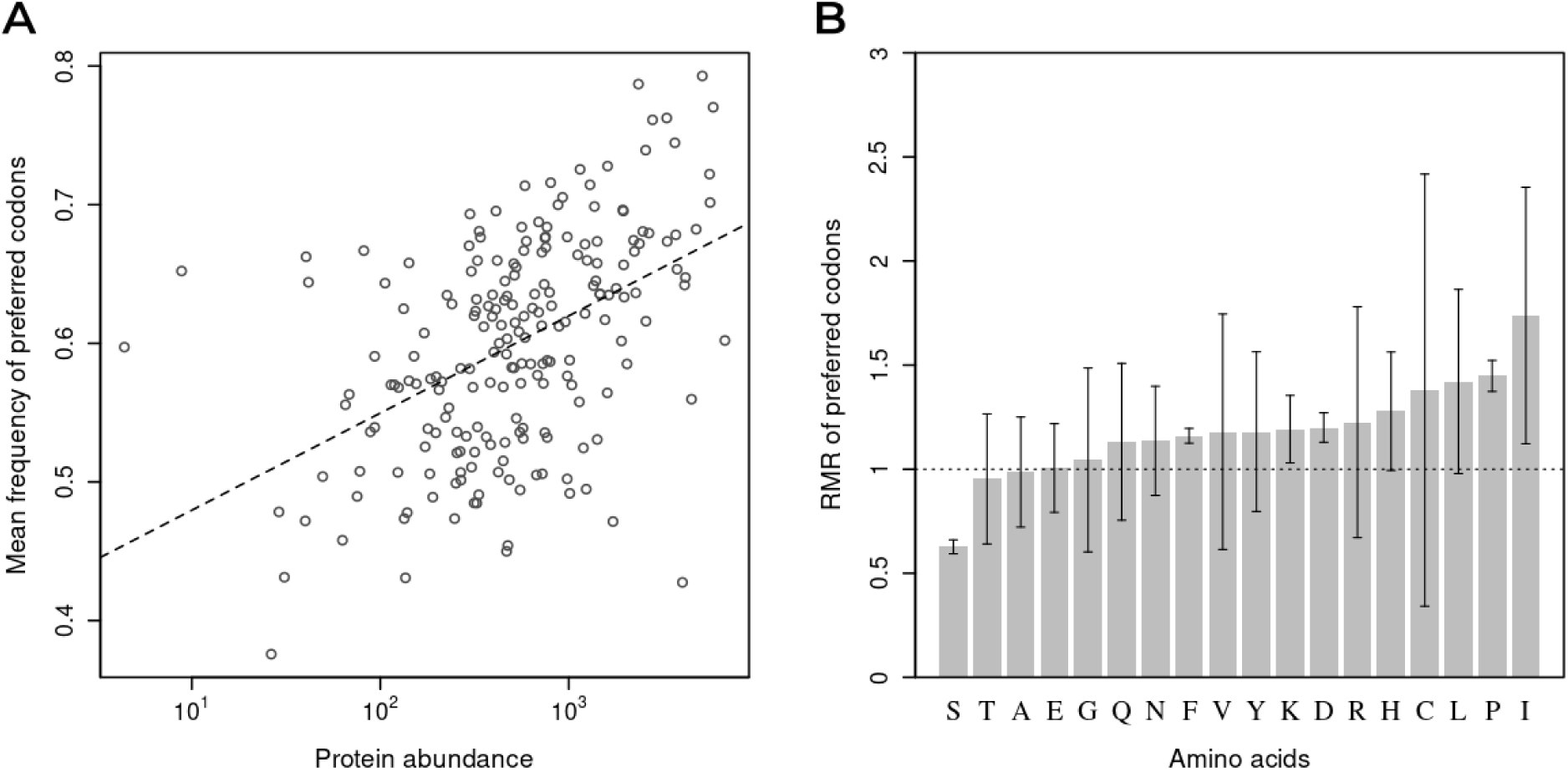
Translational accuracy of codons favored in highly expressed genes. A) Enrichment of preferred codons in highly expressed genes. The enrichment is measured as the relative frequency of preferred codons among synonymous codons in a gene averaged over amino-acids. The linear regression coefficient for a correlation between this measure of codon usage bias and log protein abundance for 205 genes (data from PaxDb) was *β* = 0.070 (SE = 0.009) and the Spearman rank correlation coefficient was *ρ* = 0.50 (*P* = 3.51 × 10^-14^). B) Relative mistranslation rate (RMR) of preferred codons, for the 18 amino acids with more than 1 synonymous codon. RMR > 1 indicates higher error rates for preferred codons compared to the mean error rate of their respective synonymous codon groups. The height of each bar corresponds to the mean RMR of each preferred codon over three biological replicates. Error bars denote the corresponding standard errors of the mean.

With the codon-level mistranslation rates estimated in this study (Figure S9), we can directly test whether codons preferred under translational selection make fewer translation errors. We compared mistranslation rates of preferred codons with that of their non-preferred synonymous codons. For each codon of an amino acid, we calculated a relative mistranslation rate (RMR), defined as the ratio of the mistranslation rates for a codon and the mean mistranslation rate of all synonymous codons for that same amino acid^34^. RMR greater than 1 indicates that a codon is less accurate than an average codon for the same amino acid. The mean RMRs of preferred codons over three biological replicates were significantly different from 1 (two-sided Wilcoxon signed-rank test, *P* = 0.0028) (Figure 3B). However, serine was the only amino acid with a mean RMR for the preferred codon significantly less than 1. For no amino acid was the preferred codon the most accurate one. Thus, we find that codons thought to be favored by translational selection are actually more susceptible to mistranslation errors.

Translation-accuracy hypothesis of codon usage states that codons favored in highly expressed genes should be translated more accurately compared to other synonymous codons. Previous studies have found support for this hypothesis in several organisms, including *E*. *coli*, based on an indirect approach proposed by Akashi^8,34–36^. Akashi argued that mistranslation errors would be most consequential for functionally critical amino-acid residues; therefore, codons selected for translational accuracy should be enriched at evolutionarily conserved sites relative to sites that are more variable. Because we see that codons favored in highly expressed genes are translated less accurately, we expect these codons to be less frequent at conserved sites. To confirm this expectation, we applied Akashi’s test to our predicted set of preferred codons. We considered a site “conserved” if the allele frequency for some amino acid at that site was at least 98% within a collection of 232 *E*. *coli* genomes (sampled from NCBI genome database; see Methods), and “variable” otherwise. We excluded certain groups of sites known to introduce artifacts^35^, and calculated odds ratios (OR) of counts for preferred and non-preferred synonymous codons at conserved and variable sites, such that an OR > 1 indicates an enrichment of preferred codons at conserved sites (Methods). Preferred codons were not significantly enriched at conserved sites (OR = 0.98, χ^2^ = 0.60, *P* = 0.438), as expected from their high error rates. In contrast, we detected a significant enrichment (OR = 1.14, χ^2^ = 19.18, *P* = 1.2 × 10^-5^) for the alternative set of preferred codons predicted using codon frequencies of all genes (Table 2 - Column 2). All four codons in this alternative set that were distinct from our primary set of preferred codons had lower error rates (Figure S8). Thus, codons selectively-favored in highly expressed genes, being error-prone, do not show signs of selection for translational accuracy. This further supports the conclusion that translational selection is not driven by advantages associated with accuracy.

### Protein abundance drives translation inaccuracy

A key prediction of the translational accuracy hypothesis is that highly expressed genes will evolve to be translated more accurately due to the negative fitness effects associated with mistranslating a highly abundant protein^8,37^. In contrast, we have seen in the previous section that codons enriched in highly expressed genes are translated inaccurately, leading to the expectation that highly-expressed genes will be more error-prone. To further explore this possibility, we measured the error-rate of a protein as the proportion of mistranslated amino acids among all amino-acid sequences that mapped to the protein in the mass-spectrometry data, and tested whether proteins mistranslation rates were correlated with protein abundance. Contrary to the prediction of translation accuracy hypothesis, we found that protein-level mistranslation rates are positively correlated with protein abundance (Figure 4A).

**Figure 4.**
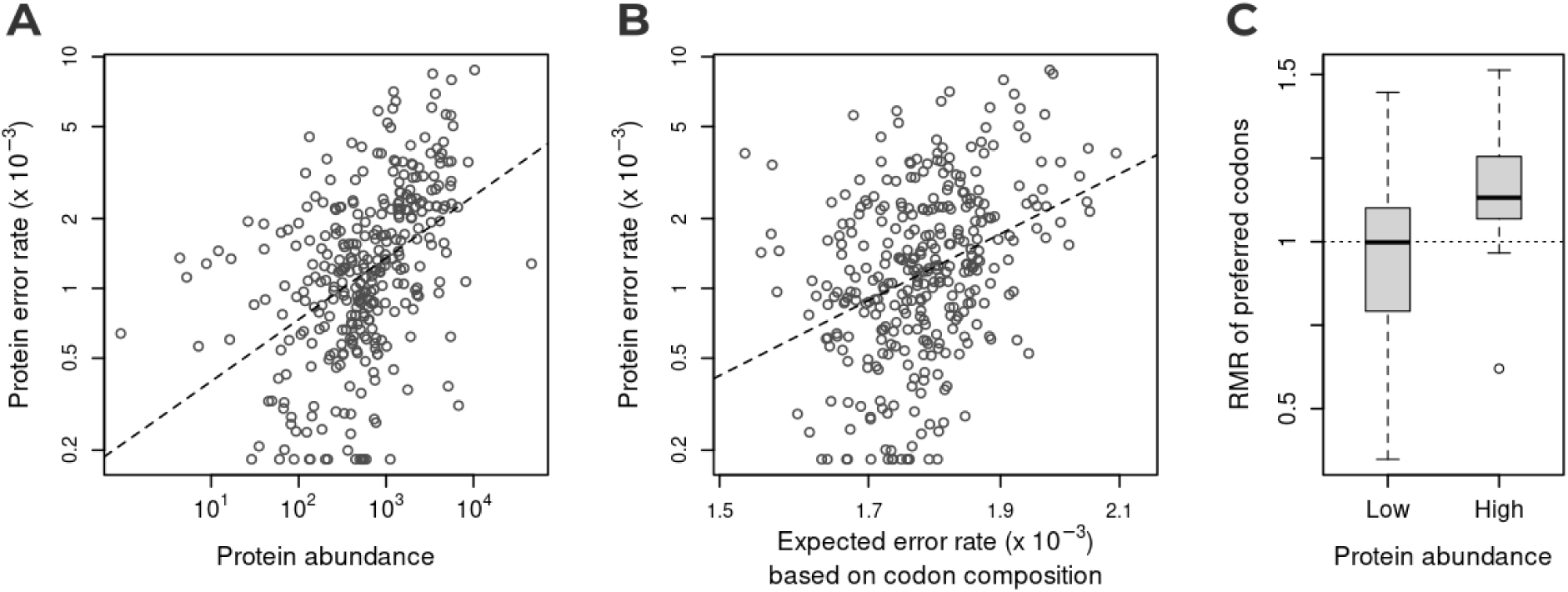
Protein abundance and codon composition as predictors of protein mistranslation rates. A) Observed error-rates of proteins are positively correlated with protein abundance. Linear regression coefficient, *β* = 0.27, *R*^2^ = 0.20, *P* = 9.1 × 10^-18^. This analysis was performed with 332 proteins with protein abundance data (PaxDB) and at least 3,000 observed codons in the mass-spectrometry data. B) Observed protein error rates are correlated with the expected error rates based on coding sequence. Linear regression coefficient, *β* = 5.96, *R*^2^ = 0.13, *P* = 1.5 × 10^-11^. C) Estimates of Relative Mistranslation Rates (RMR) of preferred codons (Table 2 – Column 1) differ between expression levels. Low and High represents the bottom 50% and top 50% of 1,714 genes respectively based on protein abundance data. Boxplots are drawn over mean RMRs of preferred codons for 18 amino acids. Outliers that are farther than 1.5 times the interquartile range outside the box are shown separately. The dashed horizontal line indicates the expected RMR of preferred codons if these did not differ from other synonymous codons in translation accuracy.

It is not evident from this correlation between protein-level mistranslation rates and protein abundance whether translation accuracy of a gene is governed by its sequence or by the level of expression. To address this, we defined an expected error rate for a protein based on its codon sequence as the sum of the error rate estimates for each codon divided by the number of codons. Observed protein error rates showed a weak but significant positive correlation with the error rates expected based on codon composition (*R*^2^ = 0.13, *P* = 1.45 × 10^-11^) (Figure 4B). A multiple linear regression of observed protein error rates on protein abundance and the expected error rates as independent variables slightly improved predictive power (*R*^2^ = 0.23, *P* < 0.001). Both predictors were significant (*P* < 0.001) but abundance was a better predictor according to the standardized coefficients of the regression model (0.36 vs. 0.20).

This dependence of the mistranslation rates of proteins on their expression could bias the estimates of codon-specific error rates. Because preferred codons are enriched in genes with high expression, which are also error-prone, the estimated error rates of preferred codons could be spuriously upwardly biased owing to this secondary effect. To test for this possibility, we estimated codon-level error rates separately for high-expression (top 50%) and low-expression (bottom 50%) set of genes. Indeed, we found that the RMR for preferred codons was higher in highly expressed genes than in lowly expressed genes (one-sided Wilcoxon signed-rank test, *P* = 0.013) (Figure 4C). However, the RMRs of preferred codons even in the low-expression set was not significantly different from 1 (two-sided Wilcoxon signed-rank test, *P* = 0.349). Thus, we find that preferred codons do not differ from other codons in their accuracy, when their enrichment in highly expressed genes is controlled for. Taken together, these results suggest that the differences in translational accuracy of synonymous codons have a minor effect on the mistranslation rates of genes compared to gene expression, and are not responsible for stronger codon bias in highly expressed genes.

## Discussion

We have presented a method that enables the determination of both the rate and spectra of translation errors. Making use of historically characterized ribosomal variants of *E*. *coli* to verify the method, we had expected translation-error rates to reflect their originally assigned phenotypes: wild-type, accurate, and error-prone^2,11,17^. However, when considering nearly the entire spectrum of possible mistranslation events, as opposed to a single codon in a single protein, the overall error rates of these variants did not significantly differ from one another. What did change were the types of mistranslation events that were enriched in each variant (Tables S2–S5). The differences among the mistranslation profiles were likely related to either kinetic or structural differences specific to each ribosome. The “error-prone” ribosomal variant struggled to accurately discriminate near-cognate codons that differed in the 3^rd^ position, which is consistent with previously reported kinetics suggesting that the error-prone variant fails to reject near-cognate tRNA at a rate approximately 10-fold higher than wild-type^2^.

The method presented here enables evaluation of nearly the entire spectrum of amino-acid substitutions, revealing which mistranslation events are most significantly enriched. For wild-type, the most significantly enriched substitution was alanine to serine. Alanine to serine substitutions have been previously shown to be caused by tRNA mischarging^5,38^. The alanine tRNA synthetase is unique in that it can mischarge with amino acids that are both larger and smaller than its cognate amino acid, and as such there is also enrichment for the other common Ala mischarging event, Ala to Gly. In fact, alanine tRNA mischarging is so pervasive across the Tree of Life that additional genomic elements have evolved to proofread charged alanine tRNA^6,39^. Despite this, the previously estimated rate of mischarging by the alanine synthetase in *E*. *coli* of 2.00 × 10^-3^ is approximately equal to the overall translation-error rates observed here for the wild-type and accurate ribosomal variants^5,6,38^. Additionally, only a single nucleotide needs to be misread for alanine to be substituted by either serine or glycine. This combination of high mischarging rates and short mutational distance leads to alanine mistranslation events to predominantly result in the misincorporation of serine and glycine.

This pattern holds true for the second most enriched substitution, Phe to Tyr, as well^5,40^. Highly enriched Trp to Phe substitutions might be explained by the steric compatibility of their synthetases^41^. For other highly enriched substitutions, like, Thr to Asp (2 MMD) and Phe to Glu (3 MMD), there is no prior literature of this mistranslation being accounted for by tRNA misactivation, but several other enriched substitutions among the top 5% of the Z-score values have been previously reported to be caused by tRNA mischarging^5,6,38,40–44^.

Considering the variation in error spectra across ribosomal variants, the “accurate” variant differed from the wild-type in only two substitutions – Gln to Leu/Ile and Asp to Asn, whereas the “error-prone” variant exhibited seven differentially enriched substitutions. Six of those seven involved substitutions with a minimum mutational distance of 1 nt, and four of those six occurred in the third position of the codon, again illustrating that the error-prone variant commonly misreads near-cognate codons.

The accurate and error-prone *E*. *coli* ribosomal variants were both classified as such via the results of a reporter assay with a single nucleotide change at the third position^2^. AAU Asn is used as the experimental codon in luciferase assays because it is the near-cognate codon of Lys AAA, the wild-type amino acid in the active site of luciferase^2,7^. However, in the present study, it was found that the error rate at AAU codons was approximately twice that of the other ribosomal variants. Additionally, mistranslation at AAU codons in the “error-prone” variant resulted in lysine misincorporation 52% of the time, whereas the other ribosomal variants never incorporated lysine at AAU codons (0%). As such, the “error-prone” ribosomal variant may have been misclassified as a consequence of the previous experimental design.

Codon bias has long been thought to relate to either translation accuracy or translation efficiency^45–48^. Lacking direct measurements of translation accuracy, others have attempted to link synonymous codon usage to translational accuracy through applications of Akashi’s test of enrichment of preferred codons at conserved amino-acid sites^8,35^. With direct measures of codon-level mistranslation rates obtained here, we show that codons selectively favored in highly abundant proteins are not translated more accurately and, accordingly, are not enriched at conserved sites. Stoletzki and Eyre-Walker (2007) observed a significant enrichment at conserved sites of preferred codons identified by Sharp and Li (1987), but we did not observe such an association for this set of preferred codons either (Figure S10). These differences could be due to their use of three ecologically distinct strains of *E*. *coli*, where it is difficult to distinguish sites that are less conserved due to weaker functional constraints from sites under divergent selection. In contrast, we analyzed hundreds of *E*. *coli* isolates, with an intermediate-level of divergence and evidence of ongoing gene-flow, from which sites accumulating neutral variation should be more easily detectable.

We find that protein-specific translation-error rates are positively correlated with protein abundance (*R*^2^ = 0.20, *P* < 0.001). This finding is somewhat unexpected given prior hypotheses that abundant proteins should be translated more accurately^8,37^. However, these prior studies have lacked direct evidence of translational accuracy; instead, accuracy was indirectly assessed using simulations to determine whether translation errors led to peptide misfolding or aggregation^8,37^. What makes a highly expressed protein more error-prone? Translational speed and accuracy tradeoffs have been studied in great detail, but the general thought has been that ribosomes are optimized for speed rather than accuracy^49–51^. We have shown here that certain codons are enriched in highly expressed genes beyond the frequencies expected based on mutational biases alone. Almost all of these “preferred codons” (14 of 16) use either the most abundant tRNA, or end in a G or C, which is known to be decoded more quickly (exceptions being Val and Arg)^52^. Numerous studies have concluded that abundant tRNAs enhance translational speed, and conversely that rare codons slow translation as the ribosome awaits the correct tRNA^48,52–60^. The codon-decoding rate directly affects ribosomal elongation and protein folding, as even synonymous single-nucleotide polymorphisms can result in alternative protein folds^61,62^. Consequently, rare codons typically occur near protein-domain boundaries, plausibly because pausing enhances the co-translational folding of those structures^62–65^. Others have also theorized that highly abundant proteins are more tolerant of missense errors; that is, they fold properly despite mistranslation^66,67^. However, a ribosomal-profiling study concluded that codons associated with rare tRNAs did not slow translation, and ribosomal pausing was instead affected by Shine-Dalgarno-like features in the mRNA^68^. Relatedly, two arginine codons are actually the least used codons in *E*. *coli* and those codon compositions (AGG, AGA) in tandem could mimic a Shine-Dalgarno-like feature^57^. Other recent ribosome-profiling studies have also, perhaps prematurely, concluded that mRNA actually does not play a role in translational speed^69,70^. Slowing of translational speed was instead reported to be caused by the translation of positively charged amino acids; primarily Arg and Lys, and to a lesser extent, His^69^. Thus, it is clear that the rate of translation is modulated by many different factors, such as: tRNA abundance, codon composition, mRNA-level features such as Shine-Dalgarno-like sites, or secondary structure; positively charged amino acids and repeated amino acids. Preferred codon enrichment among highly abundant proteins may be driven by all of these factors to enhance translational speed, or additionally enable robust folding of highly expressed proteins.

Based on the data obtained in this study, we posit that for highly expressed proteins, selection favors speed of translation over accuracy. Moreover, among highly expressed proteins, mistranslation events might be well-tolerated as an abundance of protein might overcome any fitness deficits caused by a single, or even a few, non-functional proteins among the population. In contrast, for proteins that are produced in low amounts, accurate translation may be more important as any defects would affect a proportionally larger fraction of the total. Indeed, the notion that translation prioritizes speed rather than accuracy has support via biophysical models, given that the level of error experienced is tolerable^25,51^. Additionally, one kinetic model described that error-rate minimization occurs at low speeds, consistent with Hopfield and Ninio^71,72^ and with the data obtained here that less abundant proteins containing more slowly decoded codons are more accurately translated^51^. Synonymous codons do differ in their translation accuracy but the enrichment of certain codons in highly expressed genes, albeit driven by natural selection, has not resulted in more accurate translation.

## Acknowledgements

This work was supported by National Institutes of Health, 2R35GM122566 and the National Science Foundation, DBI-2119963. The authors acknowledge resources and support from the Knowledge Enterprise Biosciences Core Facilities at Arizona State University. We would like to acknowledge Tim Karr and Jessica Sandler from the Knowledge Enterprise Biosciences Core Facilities at Arizona State University for mass spectrometry related sample processing and instrument operation.

## Author Contributions

Conceptualization, R.S. and M.L., methodology, R.S., F.A., and W. -C. H., software, R.S., F.A., and W. -C. H. formal analysis, R.S., F.A., and W. -C. H., investigation, R.S., F.A., and T.L., writing – review and editing R.S., F.A., W. -C. H., T.L., and M.L. funding acquisition, M.L., visualization, R.S. and F.A., supervision, R.S. and M.L.

## Data and Code Availability

The method presented here for directly searching mass spectrometry data for error-containing peptides is amenable for use with existing mass spectrometry data. The code to perform this analysis is available on GitHub (github.com/LynchLab/ErrorRateAnalysis). The mass spectrometry proteomics data have been deposited to the ProteomeXchange Consortium via the PRIDE^73^ partner repository with the dataset identifier PXD064116 and project DOI of 10.6019/PXD064116.

## Methods

### Culture conditions

*E*. *coli* ribosomal variants (a generous gift from Dr. Hani Zaher) were grown overnight at 37°C in Luria-Bertani broth (Miller formulation) with aeration. Cultures were grown to stationary phase prior to protein purification.

### Protein Purification and Peptide Preparation

Cells were pelleted via centrifugation and frozen prior to lysis. Lysis was achieved by incubating the pellets in 25 µL pH 7.5 lysis buffer (pH adjusted with phosphoric acid) consisting of 5% sodium dodecyl sulfate (SDS) and 50 mM triethylammonium bicarbonate (TEAB) at 95°C for 10 minutes. Samples were kept on ice unless otherwise stated. Samples were then clarified by centrifugation for 10 minutes (13,000 RCF). Proteins (the supernatant) were reduced via the addition of dithiothreitol (DTT) to a concentration of 50 mM with a 10-minute incubation at 95°C. Iodoacetamide (IAA) was added to a concentration of 40 mM, and samples were incubated at room temperature, in the dark, for 30 minutes to irreversibly modify cysteines to carbamidomethyl cysteine, preventing them from forming disulfide bonds. Phosphoric acid was added to a final concentration of 5.5%, then 2 µL of 1 µg/µL trypsin was added to the acidified samples. S-trap binding buffer (100 mM TEAB in 90% methanol, pH 7.1) was then added to the peptide samples (volume equal to 6 times the sum total of DTT, IAA, and phosphoric acid). Samples were then transferred to S-trap columns and centrifuged at 4,000 RCF for 1 minute. The column was washed four times with 150 µL of the S-trap binding buffer. On-column trypsin digestion was performed by adding 25 µL of 50 mM TEAB, pH 8, containing 0.5 µg of trypsin to the top of the protein trap. The column was then incubated for 1.5 hours at 47°C. Peptides were finally eluted three times (centrifugation of 4000 RCF for 1 minute), first with 40 µL of 50 mM, pH 8, TEAB, then with 0.2% aqueous formic acid, and then with 35 µL of 50% acetonitrile containing 0.2% formic acid. Peptides were dried in a speed vacuum for two hours, then resuspended in 20-40 µL of 0.1% formic acid.

### Mass Spectrometer Settings

Approximately 500 ng of peptides were injected onto an Ultimate 3000 HPLC (Thermo Fisher Scientific) at 0.300 nL/min and separated on a 50 cm EASY SPRAY C18 column (Thermo Fisher Scientific) before being analyzed in an Orbitrap Fusion Lumos Tribrid Mass Spectrometer (Thermo Fisher Scientific). Buffer A consisted of 99.9% LC-MS-grade water, Buffer B consisted of 99.9% LC-MS-grade acetonitrile, and both buffers contained 0.1% formic acid. A 5 minute equilibration preceded a 60 minute analytical gradient of 1%-35% B and a 5 minute column wash at 80% B. The mass spectrometer was run in database-dependent acquisition (DDA) mode with both MS1 and MS2 being detected on the Orbitrap. The MS1 full scan resolution was set to 120K, a scan range of 375-1500 m/z, charge states were set to the range of +2 to +7, the AGC target was set to 400K, and maximum injection time was 50ms. Precursor ions were activated by HCD with 30% collision energy, MS2 resolution was set to 30K, the AGC target was set to 50K, and the maximum injection time was 54ms. Raw data files were analyzed in Proteome Discoverer 3.1 (Thermo Fisher Scientific).

### Whole-Genome Sequencing and Generation of Annotation Files

The ribosomal variants of *E*. *coli* were subjected to whole-genome sequencing. Specifically, hybrid sequencing (a combination of Illumina and Nanopore sequencing) which was carried out by SeqCoast Genomics. DNA was extracted using a MagMax Microbiome Ultra Nucleic Acid Isolation kit. For Illumina sequencing, samples were prepared for using the Illumina DNA Prep tagmentation kit and IDT For Illumina Unique Dual Indexes. Sequencing was performed on the Illumina NextSeq 2000 platform using a 300-cycle flow cell kit to produce 2×150bp paired reads. 1-2% PhiX control was spiked into the run to support optimal base calling. Read demultiplexing, read trimming, and run analytics were performed using DRAGEN v4.2.7, an on-board analysis software on the NextSeq 2000. For Nanopore sequencing, DNA samples were prepared using the Oxford Nanopore Technologies SQK-NBD114 native barcoding kit. Long Fragment Buffer was used to promote longer read lengths. Sequencing was performed on the GridION platform using a FLOW-MIN114 Spot-ON Flow Cell, R10 version with a translocation speed of 400bps. Base calling was performed on the GridION using the super-accurate base calling model with barcode trimming enabled. Raw Illumina reads were trimmed using Trimmomatic software (v0.39), on default settings^74^. Raw Nanopore reads were trimmed using Porechop (v0.2.4)^75^. High-quality reads were assembled using the Unicycler software (v0.4.4) and annotated using BAKTA (v.1.5.1)^76,77^.

### Initial and Custom Database Search

FASTA files for gene sequences and protein sequences were generated from genome annotation files derived from whole-genome sequencing. These annotation files were processed with a custom parser that outputs FASTA files for validated protein and DNA sequences. Resultant protein FASTA files were loaded into Proteome Discoverer 3.1 as the protein database. The initial database search in Proteome Discoverer utilized the Sequest HT node for database searching, the wild-type proteome for each ribosomal variant, and the Percolator node for PSM validation^20^. In Sequest HT, trypsin (full) was selected for proteolytic digestion with up to 2 missed cleavage sites permitted, a precursor mass tolerance of 10 ppm, and a fragment mass tolerance of 0.02 Da. Dynamic modifications included oxidation and protein N-terminal acetylation; static modifications included only carbamidomethylation of cysteines. Default settings were used in Percolator to calculate both posterior error probabilities (PEP) and q-values to a target FDR of 1%^20,78^. The custom database (see Custom Database Generation) search was carried out with the same settings with the only difference being that the custom database containing the mutant peptides was loaded instead of the variant-specific wild-type proteome. Exported data of the custom database search utilized Percolator-derived PEP values for PSM validation (0.01 PEP), as PEP is statistically more conservative than FDR, and is better suited for the validation of individual PSMs^21^. Those exported data were subjected to analysis with a custom python script (see Translation Error Rate Data Analysis).

### Custom Database Generation

The “Proteins” tab was exported from Proteome Discoverer and processed through a custom python script that results in the *in silico* trypsinization of detected proteins followed by an internal homology check, then all possible amino acid substitutions were made to peptides that passed the internal homology check, as well as length and mass filters. Length was set to six amino acids and mass was set to 500 Da; additionally, peptides containing selenocysteine were filtered out. The *in silico* trypsinization was set to allow for up to two missed cleavages due to inefficiency in the trypsin enzyme. The number of missed cleavages can be adjusted to any number, though two is sufficient. Additionally, the number of substitutions to make in each peptide should only be set to one; all analysis scripts here have only been written to accommodate single substitutions. Moreover, substitutions can be set to zero if only trypsinized products for a given missed cleavage number is desired. Substitutions can be set to two or three as well, but is computationally prohibitive. The internal homology script filters out trypsinized peptides from different proteins that differ by a single amino acid in their wild-type sequences. If two peptides were identical, then it checks the DNA sequence of both peptides, and if the DNA is also identical then the peptide is kept. In this way, the amino acid and codon level error rates are unaffected, and thereby the overall translation error rate is unaffected as well. This becomes relevant when determining protein-level error rates and protein abundance relationships, as such, those peptides are removed from analysis at that step. In sum, only proteins that were detected in the initial search were subjected to *in silico* trypsinization, an internal homology check, and mutagenesis. The resultant FASTA file contains both the wild-type parent peptides and the systematically mutagenized daughter peptides.

### Translation Error Rate Data Analysis

After performing the Proteome Discoverer search with the custom database, the analysis result tab of interest within the program was “Peptide Groups.” This tab of data was exported and then processed through a custom python pipeline. The information gathered from the exported data were the sequence column, # PSMs column, and the PEP column. These detected sequences were filtered on PEP scores with a cutoff of 0.01, any value greater than the cutoff was filtered out. A 1% posterior error probability (PEP) cutoff was utilized because PEP is better suited for validation of single PSMs, whereas FDR is better suited for the identification of proteins given many PSMs^21^. After loading the custom database, peptide sequences containing Ile to Leu or Leu to Ile substitutions were searched for and reverted to wild-type, as Leu and Ile are indistinguishable from one another within the conditions of the mass spectrometry performed here. This would result in a small amount of downward bias of the measured translation error rate. Detected sequences were then matched to the sequences contained in the custom database file. Some detected sequences did not have an exact sequence match because they did not conform to the predicted trypsin digestion fragmentation. This was caused by either the *in vivo* mistranslation events of the trypsin recognition amino acids (K, R, or P), or via collision induced dissociation within the mass spectrometer. In all likelihood, those peptides named non-standard products (NSPs) were derived from mistranslation events of the trypsin related amino acids. Proteome Discoverer performs *in silico* enzymatic digestion on-the-fly during the search process because it treats each pre-trypsinized peptide as a full-length protein. Therefore, the supplied mutant database file included all possible mistranslation events that would affect trypsin cleavage; Proteome Discoverer generated those products and searched for them. Not all newly generated trypsin sites will result in two detectable peptides, as the location of the trypsin site can result in an asymmetrical cleavage that results in a peptide that falls below detection limits. Detected NSPs are then mapped back to their parent peptides during the execution of the pipeline. An internal homology filter for NSPs is also implemented to exclude NSPs that also happened to map to an internally homologous protein region. Once all detected sequences have been mapped to their peptide identifiers, which the peptide identifier includes the type of substitution made, then peptides belonging to a set of specific substitutions were filtered out due to chemical artifacts. The table of artifacts is shown below in Table 3:

**Table 3.** Chemical modifications designated as artifacts.

| Substitution | Monoisotopic Mass | Average Mass | Mass Shift | Modification |
| --- | --- | --- | --- | --- |
| Asn → Asp | 0.984016 | 0.9848 | 1 | Deamidation |
| Gln → Glu | 0.984016 | 0.9848 | 1 | Deamidation |
| Glu → Ser | -42.010565 | -42.0367 | -42 | Unknown |
| Ser → Asp | 27.994915 | 28.0101 | 28 | Formylation |
| Thr → Glu | 27.994915 | 28.0101 | 28 | Formylation |
| Ser → Ala | -15.994915 | -15.9994 | -16 | Deoxidation |
| Tyr → Phe | -15.994915 | -15.9994 | -16 | Deoxidation |

Due to identical mass deltas, these substitutions are indistinguishable from chemical modifications that may occur either naturally, or through sample preparation for mass spectrometry.

Values shown in Table 3 were obtained from Unimod, a database of protein modifications for mass spectrometry^79^. Among the artifacts listed here is Glu to Ser, which did not have a modification within 0.02 Da of the substitution mass delta listed in Unimod, though the observed data highly suggest that this mass delta is being caused chemically. For reference, when including that substitution in the data analysis, Glu experienced an unusually high error rate of 1.5% with 81% of substitutions mapping to Ser. For comparison, when Glu to Ser is treated as an artifact (removed from analysis), then the measured translation error rate of Glu becomes realistic at approximately 3.00 × 10^-3^. Lastly, counts of all wild-type amino acids and all mutant amino acids were taken, then the error rate was calculated as the number of mutant amino acids divided by the sum total of amino acid sites sampled.

### Generation of Substitution Spectra

Both amino acids and codon-level substitution spectra were generated during the execution of the custom pipeline. Each mutant peptide has an identifier associated with it that denotes the wild-type amino acid and the substituted amino acid. Provided each identifier and the number of peptide spectral matches (PSMs) associated with each peptide sequence, a number of specific substitutions was determined for each amino acid pair in a 20 × 19 matrix, wherein 19 represents the destination amino acids where Ile and Leu have been combined into a single Xle row. This matrix represents the substitution spectra associated with detected mistranslation events. Substitutions identified as artifacts, due to indistinguishable mass differences between certain amino-acid substitutions and chemical modifications, were excluded from the analysis. The error rate per amino acid was calculated by taking the number of observed mistranslation events per amino acid and dividing it by the total number of codons sampled for that amino acid. Together, these matrices report the likelihood of a mistranslation event per amino acid, and the bias towards the incorporation of any other amino acid given a mistranslation event. The codon substitution spectrum was determined in a way very similar to the amino acid substitution spectrum. In any detected mistranslated peptide, the exact DNA coordinates were determined, and therefore the codon that was mistranslated was determined. This allowed for the generation of a matrix that would reveal any bias in specific codons towards specific amino acids.

### Identification of significantly enriched substitutions

To determine which types of translation errors occurred more frequently than expected by chance, we performed simulations to generate a randomized dataset for comparison. 10,000 rounds of simulation were performed to generate pseudo datasets based on the marginal distributions of amino acid sources and destinations. The marginal probability of an amino acid serving as a source or destination was defined by the proportion of the total counts per the amino acid source or destination divided by the total number of observed mistranslation events (n_wild-type_ = 6,692). In each round of simulation, pairs of source and destination amino acids were drawn with the function *sample* in R according to the marginal probabilities, but those deemed as artifacts (E-to-S, N-to-D, Q-to-E, S-to-D, S-to-A, T-to-E, and Y-to-F), or those with the same amino acid (including I-to-X and L-to-X) were discarded. The drawing and discarding step continued until a total of 6,692 pairs were acquired, which generated a pseudo dataset for a simulation round. After performing 10,000 rounds of simulation, for each substitution, the nominal *P*-value was determined by the proportion of pseudo datasets where the simulated observed numbers were larger than or equal to the empirically observed number. Given that only 10,000 rounds of simulations were performed, the case where no simulated observed value was larger than or equal to the empirically observed number was noted as *P* < 0.001. Under Bonferroni‘s multiple testing correction (353 comparisons) and 5% family-wise error rate, only the nominal *P*-values smaller or equal to 0.001 were considered statistically significant. Using 10,000 simulated numbers per substitution, the mean and standard deviation were calculated. The Z-score for a substitution was then determined by the difference between the simulated mean and the empirical value, normalized by the simulated standard deviation.

### Abundance Analysis

Protein abundance data were obtained from PaxDb integrated datasets^28^. These datasets are downloaded with non-universal protein identifiers, which requires some ID linking between parsed proteomes obtained from the custom parsing program used here. The custom parser results in proteomes with unique identifiers, such as locus tags or protein IDs. As such, a custom script was made that links PaxDb identifiers to the IDs found in the parsed proteome file by comparing protein sequences found in local files, or by fetching protein sequences from NCBI. The custom script optionally accepts a proteome file that utilizes the PaxDb identifiers as FASTA headers. In this case, the PaxDb identifiers were from the STRING database^80^, so the *E*. *coli* reference proteome was downloaded from STRING. The custom script was executed with the STRING reference proteome and the proteome generated from the custom parser. Protein sequences in these two files were compared, if they were found to be identical then the parsed ID was linked to that PaxDb identifier. Following assignment of exact matches, PaxDb IDs that still lacked a linkage to the parsed proteome were subjected to pairwise alignment of all other protein sequences in the parsed proteome. If the alignments yielded a single match with greater than 95% identity, then the parsed ID would be linked to the PaxDb ID. Alignments, that fell between 90 and 95% identity were manually reviewed. Following the local search, STRING IDs were converted to gene symbols by retrieving the preferred names from STRING; preferred names were confirmed or revised using a public gene annotation web-service, MyGene.info^81^. These gene symbols were used to programmatically retrieve protein sequences from NCBI. Protein sequence retrieval from NCBI first uses gene symbols obtained from MyGene.info, then if any IDs remained unlinked, PaxDb preferred names are then used. These sequences were used as described before, by first searching for exact matches among the annotated protein sequences, followed by alignments. If a gene symbol listed in the PaxDb database remains unlinked, it is assumed that the corresponding protein was either not present in the annotation file or did not meet the minimum identity threshold of 90%. The vast majority of proteins had sequence identity matches greater than 98%. Once all possible links have been made between the annotated proteome and the abundance data, then a custom script is executed for abundance analysis. This script finds all peptides that belong to a single protein in the analyzed data, then counts all wild-type and mutant amino acids for that protein. Peptides that could map to multiple proteins are removed from the analysis. The error rate of the protein is determined by taking the sum total of amino-acid substitutions for that protein, then dividing that value by the total number of amino acids sampled for that protein. Only proteins that met a minimum requirement of 3,000 sampled amino acids were considered for further analysis.

### Identification of Orthologs

To measure codon frequencies in the *E*. *coli* genome and to identify conserved and variable amino acid sites for Akashi’s test of translation accuracy, orthologs of the proteins annotated in the wild-type strain were identified in a collection of 232 other *E*. *coli* genomes, including the K-12 MG1655 reference strain. These genomes were selected from a random sample of 1,000 genomes from NCBI after filtering out highly similar genomes, using PopPUNK^82^ and lineages that appear genetically disconnected with the rest of the population, using PopCOGenT^83^. The ortholog search was performed using a reciprocal best-hit approach with BLASTp^84^, excluding hits with less than 80% query aligned and log2-fold difference in subject and query length exceeding 0.5. Protein accessions for the orthologs of the wild-type strain identified across this collection of *E*. *coli* isolates are provided in Supplementary Data File 1. Amino-acid alignments were generated using MUSCLE (v3.8.1551) and codon alignments were generated using PAL2NAL (v14)^85,86^. For the proteins present in at least 220 out of 233 strains (∼ 95%) *i*.*e*., “core” proteins, frequencies of 61 codons at each site, excluding the first 50 and the last 20 sites, were determined.

### Identification of Preferred Codons

Preferred synonymous codons for different amino acids in *E*. *coli* were identified based on population-genetic analysis of codon frequencies. The Li-Bulmer model of mutation-selection balance was applied to the observed frequencies of codons at fixed sites while accounting for the effect of mutation biases on codon usage bias^29,30^. Codon sites were considered fixed if the most frequent codon was present in at least 98% of the sequences, based on the multiple sequence alignments of protein-coding genes generated above. The relative mutation rates among 4 nucleotides were derived from the mutational spectrum of the *E*. *coli* strain PFM2 wild type obtained through a mutation accumulation experiment^31^.

The original Li-Bulmer model provides an expression for the equilibrium probability of one of the two alleles under mutation-selection balance, *p* = 1/(1+***β***e^-S^) where *S = 2 N*_e_ *s* with *N*_e_ being the effective population size and *s* being the selection coefficient for the target allele, and ***β*** is the mutation bias towards the other allele. A 4-allele extension of this model was applied to the frequencies of four codons in a codon group (defined by the first two codon position). The construction of this model can be understood by noting that the above equation represents a solution to the stationary distribution of a 2-state Markov chain with transition matrix, *P* = 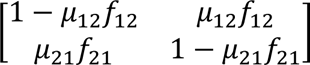 where *μ*_ij_ is the mutation rate from allele *i* to allele *j* and *f_ij_* = 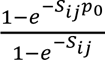 is the fixation probability of *j* over *i* and *S_ij_* = 2 *N_e_s*_ij_ is the corresponding population-scaled selective advantage, with *p*_0_ being the initial population frequency of allele *j*. The stationary distribution, π = (*p*,*q*) that satisfies π = π*P*, can be found by solving a linear system of 2 equations under the constraint that *p* + *q* = 1^87^. The solution for the above matrix is 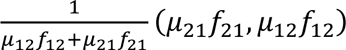. The Li-Bulmer equation can be recovered by replacing *μ*_12_/*μ*_21_ with *β*, and noting that *f*_12_/*f*_21_ equals *e*^-S^ where *S* = *S*_21_.

A 4-allele model of mutation-selection balance was derived by similarly considering the stationary distribution of a 4-state Markov chain. The 4 nucleotides were labeled in the alphabetical order *i*.*e*., 1 (A), 2 (C), 3(G) & 4(T), and mutation rates (*μ*_ij_) were set to the six base-pair substitution rates derived from the mutation-accumulation study *eg*., *μ*_24_ = *μ*_31_ = *µ_GC→AT_*. The selection coefficient for nucleotide *A* was set to 0 and the other 3 selection coefficients (*S*_j_) were relative to A, so that the selection term (*S*_ij_) in fixation probability of allele *j* over *i* was set to:

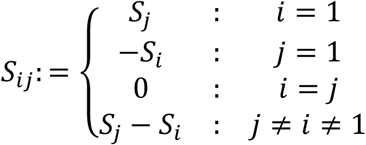

Expressions for equilibrium probabilities as functions of 3 selection coefficients were derived by solving the resultant system of linear equations. Values of selection coefficients were derived from these equations based on the Newton-Raphson method^87^. The Newton-Raphson approach finds roots of the vector equation **f**(**x**) = **0** based on iterations of the form **x**_t+1_ = **x**_t_ – (*D***f**(**x**_t_))^-1^ **f**(**x**_t_) where *D***f**(**x**) is the Jacobian matrix. In our problem, **f**(**x**) represents the difference between equilibrium and observed frequencies of the 4 codons in a group. Therefore, estimated selection coefficients correspond to values that equate the expected equilibrium frequencies under mutation-selection balance to their observed values. Starting estimates for selection coefficients were arbitrarily set to (0.1, 0.2, and 0.3). This analysis was performed in Mathematica 13.2. For amino acids with less than 4 synonymous codons, selection coefficients initially estimated relative to codon NNA were recalculated relative to the smallest selection coefficient among synonymous codons. Preferred codon for each amino acid was identified as the one with the highest selection coefficient. For 3 amino acids with 6 synonymous codons, *i*.*e*., codons belonging to two-distinct codon groups, the preferred codon was identified from the group with the higher selection coefficient.

### Measure of codon usage bias

The codon usage bias of a gene was quantified as the proportion of preferred codons out of the total number of codons of an amino acid in that gene, averaged over 18 amino acids with more than 1 synonymous codon. This metric was used to test for the enrichment of preferred codons in highly expressed genes. It was calculated for 1,100 genes with at least 4 occurrences of 16 out of 18 amino acids with preferred codons, and at least 30 codons left after removing the first 50 and the last 20 codons.

### Akashi’s Test of Translation Accuracy Hypothesis

The enrichment of preferred codons at sites with conserved amino acid residues was tested following the method used by Akashi 1994^36^. Conserved amino acid sites were defined as the sites where the most frequent amino acid was at least present among 98% of the 232 *E*. *coli* isolates. Non-core proteins, proteins missing from the wild-type strain, and those without an ortholog or more than one ortholog in *Salmonella enterica* were excluded. Contiguous stretches of mutations were removed, along with the first 50 and the last 20 codons of each gene. Sites with gaps, stop codons, or undetermined codons in any sequence were removed. Sites where the wild-type residues are among the 18 amino acids with preferred codons identified as above, and are the same as the most frequent amino acid at the site, were retained. 2 × 2 contingency tables of the numbers of preferred codons at conserved and variable sites in the wild-type strain were created for each of the 18 amino acids with synonymous codons in each protein. The tables with zero margin totals were excluded. The remaining tables were pooled following Mantel-Haenszel’s procedure and a χ-squared test of the enrichment of preferred codons at conserved sites was performed ^88^. The Mantel-Haenszel’s statistic, W_MH_, which is equivalent to an odds ratio for the pooled dataset, was calculated as 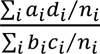 where *i* indexes a 2 × 2 contingency table corresponding to an amino acid in a gene, *a_i_*, *b_i_* stands for the number of preferred and unpreferred codons respectively at conserved sites, and *c_i_*, *d_i_* stands for the number of preferred and unpreferred codons respectively at variable sites, and *n*_i_ = *a*_i_ + *b*_i_ + *c*_i_ + *d*_i_, is the total number of sites considered for that amino acid in that gene. The χ-squared test-statistic was calculated, with a continuity correction, as 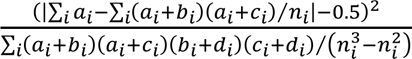. The significance of the association between preferred codons and conserved sites was evaluated under a χ-squared distribution with 1 degree of freedom.

