## Supplementary Figures for "Translation Accuracy in *E*. *coli*"

Table S1. Z-scores of observed substitutions and chi-squared values for overall error rates per amino acid of the wild-type ribosomal variant

| | R | H | K | D | E | S | T | N | Q | C | G | P | A | V | I | L | M | F | Y | W |
|---|---|---|---|---|---|---|---|---|---|---|---|---|---|---|---|---|---|---|---|---|
| R | | -1 | 1 | -2 | -2.29 | 4.43 | 0.2 | -0.96 | -1.95 | 3 | 6.3 | -2 | 0 | -2 | 1 | -1 | -0 | 1 | 7.16 | -0 |
| H | -1 | | 0 | 1 | -4.39 | -1.02 | -2.97 | 0.44 | -1.86 | -1 | -1 | -3 | -4 | 2 | 5 | 6 | -1 | -1 | 0.26 | -1 |
| K | -2 | 5 | | -3 | 4.29 | 5.77 | -0.26 | -3.7 | 9.37 | -1 | 4.9 | -3 | -2 | -2 | -2 | -3 | 3 | -3 | -1.31 | 1 |
| D | -1 | -1 | -2 | | 4.56 | | **24.47** | artifact | -4.67 | 11 | -2 | -4 | -6 | -6 | 1 | 1 | 2 | -4 | -1.53 | -1 |
| E | -2 | -1 | 2 | 0 | | -0.53 | artifact | -2.5 | artifact | -0 | -1 | 11 | -4 | -2 | -4 | -4 | 9 | **15** | 0.21 | -1 |
| S | -2 | -2 | -2 | -6 | artifact | | 5.33 | -6.27 | 2.67 | 4 | -1 | -1 | **36** | -7 | -6 | -7 | -2 | -4 | -2.21 | -0 |
| T | -2 | 3 | -2 | 2 | -5.92 | 3.91 | | 0.27 | -3.19 | -1 | 0 | 6 | -5 | **17** | -4 | -5 | -2 | -2 | -1.06 | -1 |
| N | -3 | -1 | -3 | 9 | 4.28 | 0.92 | -5.97 | | 4.93 | -2 | -1 | 5 | -8 | -6 | 1 | -0 | 4 | -1 | -0.96 | -1 |
| Q | -1 | -1 | **16** | -4 | 10.71 | 1.36 | -3.46 | 0.45 | | -2 | -0 | -4 | -4 | -2 | -2 | -2 | 2 | -1 | 0.98 | 0 |
| C | 1 | 5 | -0 | -5 | -4.1 | -1.58 | -4.72 | -5.69 | -0.41 | | -3 | -3 | -6 | -3 | 9 | **14** | -2 | -3 | 8.71 | 0 |
| G | -2 | -2 | -1 | 5 | -7.47 | 2.17 | 8.83 | 5.21 | -5.41 | -0 | | -5 | **21** | -3 | -7 | -8 | -3 | -5 | -1.77 | -1 |
| P | 2 | 3 | -3 | -3 | -4.97 | -1.6 | -2.08 | -0.12 | -4.62 | -2 | -3 | | 1 | **20** | -3 | -2 | -3 | -4 | 0.01 | 1 |
| A | -2 | -2 | -3 | -3 | 3.15 | artifact | -3.34 | 1.83 | 4.12 | -0 | **16** | -2 | | 2 | -0 | -3 | 1 | -5 | -2.09 | -1 |
| V | 3 | -2 | -3 | -4 | -9.63 | -1.71 | -1.29 | -4.08 | -6.77 | -3 | -4 | **14** | -8 | | **15** | **20** | -4 | -6 | -2.69 | -1 |
| X | 5 | -2 | 4 | **11** | -2.1 | -0.8 | -5.3 | 4.98 | 7.38 | -2 | -3 | -5 | -6 | 7 | | | -1 | -3 | -2.35 | -1 |
| M | -1 | -0 | -1 | -2 | **15.48** | -2.07 | -5.74 | **13.65** | -0.3 | -2 | -4 | -6 | -7 | -6 | -6 | -7 | | 8 | 0.9 | 0 |
| F | 1 | -1 | -1 | -1 | 0.82 | 0.73 | -2.21 | 1.16 | 6.61 | 0 | -1 | -2 | -3 | -2 | -2 | -2 | 6 | | artifact | **14** |
| Y | 2 | 0 | -1 | -3 | -2.02 | -0.86 | -1.62 | -2.99 | -1.38 | 1 | 0.3 | -3 | -3 | -2 | -1 | -4 | -1 | **26** | | 1 |
| W | 2 | -0 | 1 | -1 | -0.71 | -0.41 | -0.52 | -1.56 | -1.33 | 1 | 11 | -1 | -2 | -1 | -1 | -2 | -1 | 1 | 7.3 | |

| | | | | | | | | | | | | | | | | | | | | |
|---|---|---|---|---|---|---|---|---|---|---|---|---|---|---|---|---|---|---|---|---|
| Chi2 | 46 | 45 | 50 | 0 | **199.1** | **263.2** | 0.12 | **288.9** | 62.24 | 50 | **318** | 6 | 13 | 79 | 14 | 1 | 1 | 83 | 52.53 | 15 |

The upper matrix of values are the Z-scores determined by comparing the observed number of substitutions to the distribution of 10,000 simulated numbers of expected substitutions, dependent on marginal probabilities of sources and destinations. The top 5% of Z-scores are bolded. The lower matrix represents the chi-squared values of the translation error rates associated with each amino acid. The top 4 values are bolded.

Table S2. Mistranslation events in the top 5% of Z-scores for the accurate ribosomal variant

| Amino Acid Source | Amino Acid Destination | Z-score | Minimum Distance (nt) |
|---|---|---|---|
| A | S | 35.14 | 1 |
| F | Y | 29.47 | 1 |
| T | D | 23.31 | 2 |
| A | G | 23.07 | 1 |
| G | W | 22.10 | 2 |
| L | V | 20.17 | 1 |
| V | T | 18.84 | 2 |
| W | F | 17.22 | 2 |
| F | E | 17.14 | 3 |
| E | M | 16.78 | 1 |
| V | P | 16.39 | 2 |
| I | V | 16.21 | 1 |
| L | C | 16.08 | 2 |
| Q | X | 16.06 | 1 |
| D | X | 14.04 | 2 |
| N | M | 13.59 | 2 |
| K | Q | 13.55 | 1 |
| D | N | 12.92 | 1 |

The significance (nominal *P*-value < or = 0.0001 and top 5% Z-score of observation) was determined by comparing the observed number of substitutions to the distribution of 10,000 simulated numbers of expected substitutions, dependent on marginal probabilities of sources and destinations. The distance was defined by the minimum number of nucleotides required to be misread to cause the translation error observed.

Table S3. Mistranslation events in the top 5% of Z-scores for the error-prone ribosomal variant

| Amino Acid Source | Amino Acid Destination | Z-score | Minimum Distance (nt) |
|---|---|---|---|
| A | S | 38.74 | 1 |
| D | E | 32.59 | 1 |
| F | Y | 29.17 | 1 |
| A | G | 29.13 | 1 |
| L | V | 24.64 | 1 |
| T | D | 23.61 | 2 |
| V | T | 22.80 | 2 |
| H | Q | 21.02 | 1 |
| Y | W | 20.33 | 2 |
| I | V | 19.69 | 1 |
| N | K | 19.53 | 1 |
| E | M | 16.49 | 2 |
| V | P | 15.56 | 2 |
| P | V | 15.37 | 2 |
| L | C | 13.64 | 2 |
| E | D | 12.28 | 1 |
| K | R | 12.03 | 1 |
| E | Q | 11.90 | 1 |

The significance (nominal P-value < or = 0.0001 and top 5% Z-score of observation) was determined by comparing the observed number of substitutions to the distribution of 10,000 simulated numbers of expected substitutions, dependent on marginal probabilities of sources and destinations. The distance was defined by the minimum number of nucleotides required to be misread to cause the translation error observed.

Table S4. Z-scores of observed substitutions and chi-squared values for overall error rates per amino acid of the accurate ribosomal variant

| | R | H | K | D | E | S | T | N | Q | C | G | P | A | V | I | L | M | F | Y | W |
|---|---|---|---|---|---|---|---|---|---|---|---|---|---|---|---|---|---|---|---|---|
| R | | 0.9 | 6.8 | -2 | -2.01 | 3.81 | -0.56 | -1.63 | -1.68 | -1 | 11 | -2 | -1 | -2 | 1.6 | -1 | -0 | -1 | 4.38 | -0 |
| H | -1 | | -1 | 1.5 | -4.73 | -0.3 | -2.63 | -1.61 | -2.3 | -2 | -1 | -4 | -5 | 2.2 | 7.4 | 7.6 | -0 | -2 | -1.34 | -1 |
| K | -2 | 4.6 | | -2 | 4.03 | 4.36 | -1.84 | -3.3 | 10.94 | -2 | 3.8 | -2 | -2 | -0 | -4 | -3 | 3.5 | -3 | -1.25 | -1 |
| D | -2 | -2 | -2 | | 8.27 | | **23.31** | artifact | -5.56 | 9.9 | -2 | -5 | -6 | -6 | -0 | 0.6 | 2.9 | -5 | -1.6 | -1 |
| E | -2 | -0 | 1.7 | -2 | | 0.32 | artifact | -1.74 | artifact | 1.8 | -2 | 9.6 | -2 | -0 | -4 | -4 | 5.8 | **17** | 0.62 | 1.9 |
| S | -1 | -2 | -2 | -6 | artifact | | 8.39 | -6.38 | -1.01 | 1.6 | -2 | -1 | **35** | -7 | -7 | -8 | -3 | -3 | -2.2 | 0.6 |
| T | 0.7 | 4.1 | -3 | 0.8 | -5.95 | 0.51 | | 1.75 | -3.22 | -0 | -1 | 6.3 | -6 | **19** | -4 | -4 | -3 | -1 | 0.63 | -1 |
| N | -2 | -2 | -3 | 13 | 3.33 | 0.41 | -6.03 | | 4.5 | -3 | -1 | 6.6 | -8 | -5 | -0 | -3 | 2.1 | 2.2 | -0.64 | -1 |
| Q | -2 | -0 | **14** | -4 | 12.2 | 1.57 | -2.71 | -1.08 | | -2 | 0.6 | -4 | -5 | -1 | -1 | -3 | 0.8 | 0.6 | 2.61 | 0.1 |
| C | 1.6 | 5.3 | -2 | -4 | -3.99 | -1.04 | -4.39 | -5.54 | -1.52 | | -1 | -4 | -6 | -1 | 7.8 | **16** | -2 | -4 | 9.74 | -1 |
| G | -3 | -2 | -2 | 4.9 | -8.08 | 2.42 | 0.61 | 6.68 | -5.89 | 2.4 | | -5 | **23** | -2 | -7 | -8 | -1 | -5 | -1.44 | -1 |
| P | 5 | 1 | -3 | -3 | -4.38 | -0.98 | -1.18 | 0.13 | -4.52 | -2 | -1 | | 3.3 | **16** | -1 | -3 | -3 | -3 | -1.87 | -1 |
| A | -2 | -1 | -1 | -2 | 2.54 | artifact | -2.21 | 2.89 | 3.81 | -1 | 7.7 | -2 | | 4.8 | -1 | -2 | 0.4 | -4 | -1.84 | -1 |
| V | 3.3 | -2 | -4 | -6 | -10.1 | -2.5 | -0.28 | -3.29 | -7.19 | -3 | -3 | 12 | -10 | | **16** | **20** | -4 | -6 | -2.86 | -1 |
| X | 2 | -0 | 4.1 | **14** | -5.53 | -0.57 | -4.43 | 4.97 | **16.06** | -1 | -2 | -4 | -5 | -2 | | | 1.8 | -4 | -1.97 | -1 |
| M | -1 | -1 | 0.8 | -3 | **16.78** | 0.64 | -5.46 | **13.59** | 2.07 | -2 | -2 | -6 | -8 | -4 | -7 | -9 | | 5.2 | -0.02 | 0.3 |
| F | -1 | -1 | -0 | -2 | 1.77 | -0.86 | -1.78 | 2.11 | 3.29 | -0 | -1 | -3 | -3 | 0.7 | -3 | -3 | 9.4 | | artifact | **17** |
| Y | 0.9 | 0.2 | 0.3 | -3 | -1.78 | -0.95 | -2.2 | -3.44 | -2.33 | 3.4 | 1.8 | -3 | -4 | -2 | -1 | -3 | -1 | **29** | | -1 |
| W | 4.1 | -0 | 1.7 | -2 | -0.93 | -0.47 | -1.43 | -1.74 | -0.86 | -1 | **22** | -1 | -2 | -1 | -1 | -2 | -1 | -1 | 6.72 | |

| | | | | | | | | | | | | | | | | | | | | |
|---|---|---|---|---|---|---|---|---|---|---|---|---|---|---|---|---|---|---|---|---|
| Chi2 | 37 | 40 | 71 | 0 | **149.2** | **250.8** | 1.36 | **363.6** | 101.1 | 68 | **398** | 9 | 0.3 | 2.9 | 25 | 0.4 | 0.1 | 55 | 50.32 | 12 |

The upper matrix of values are the Z-scores determined by comparing the observed number of substitutions to the distribution of 10,000 simulated numbers of expected substitutions, dependent on marginal probabilities of sources and destinations. The top 5% of Z-scores are bolded. The lower matrix represents the chi-squared values of the translation error rates associated with each amino acid. The top 4 values are bolded.

Table S5. Z-scores of observed substitutions and chi-squared values for overall error rates per amino acid of the error-prone ribosomal variant

| | R | H | K | D | E | S | T | N | Q | C | G | P | A | V | I | L | M | F | Y | W |
|---|---|---|---|---|---|---|---|---|---|---|---|---|---|---|---|---|---|---|---|---|
| R | | 0.2 | **12** | -1 | -2.43 | 3.54 | -1.42 | -2.07 | -0.49 | -1 | 6.3 | -1 | -1 | -2 | 1 | -1 | -1 | -0 | 5.53 | -0 |
| H | 2.3 | | 0.3 | -1 | -4.55 | -0.48 | -2.36 | -3.16 | 5.16 | 1.6 | -1 | -4 | -5 | 2.5 | 5.6 | 3.2 | -2 | -0 | 0.61 | 0.6 |
| K | -2 | 0.7 | | -5 | 1.41 | 2.11 | -2.7 | **19.53** | 6.99 | -1 | 0.7 | -4 | -4 | -3 | -4 | -5 | -1 | -4 | -1.84 | 1.2 |
| D | -2 | -3 | -2 | | **12.28** | | **23.61** | artifact | -4.45 | 6.3 | 6.2 | -5 | -6 | -7 | 0.6 | 0.7 | -0 | -5 | -1.15 | -1 |
| E | -3 | -2 | 1.1 | **33** | | -2.05 | artifact | -2.93 | artifact | -1 | -1 | 4.1 | -7 | -4 | -7 | -7 | 0.8 | 5.6 | -1.48 | -0 |
| S | -2 | -3 | -2 | -7 | artifact | | 6.94 | -7.29 | 1.77 | 1.8 | -0 | -1 | **39** | -7 | -7 | -8 | -3 | -5 | -2.49 | -0 |
| T | -1 | 0.7 | -1 | -1 | -6.43 | 6.54 | | 0.02 | -3.75 | -1 | -2 | 4.6 | -5 | **23** | -5 | -3 | -3 | -2 | -1.32 | -1 |
| N | -2 | -2 | -2 | 7.6 | 1.26 | 1.89 | -5.14 | | 9.07 | -2 | -2 | 4.9 | -8 | -4 | -1 | -1 | 2.2 | -1 | -0.48 | -1 |
| Q | 0.4 | **21** | 11 | -6 | 11.9 | -0.69 | -4.2 | -2.19 | | -2 | -1 | -4 | -6 | -2 | -3 | -2 | 1.3 | -1 | 0.38 | 0.9 |
| C | 4.2 | 2.7 | -2 | -6 | -3.22 | -1.09 | -4.52 | -6.02 | 0.53 | | -2 | -3 | -6 | -2 | 10 | **14** | -1 | -4 | 8.52 | -1 |
| G | -1 | -3 | -2 | 1.1 | -7.64 | 3.61 | 4.43 | -1.82 | -5.35 | 0.5 | | -4 | **29** | -2 | -6 | -8 | -2 | -5 | -1.82 | -1 |
| P | 8.5 | 0.4 | -2 | -4 | -4.09 | -1.69 | -0.27 | 0.04 | -4.05 | -2 | -2 | | 3.5 | **16** | -1 | -4 | -2 | -4 | -0.42 | -1 |
| A | -2 | -2 | -2 | -4 | 4.23 | artifact | -2.36 | 4.01 | -0.14 | -0 | 9.5 | -1 | | 4.2 | -1 | -1 | 0.8 | -5 | -2.1 | 0.9 |
| V | -0 | -4 | -4 | -9 | -10.5 | -1.18 | -0.55 | -6.38 | -6.88 | -2 | -4 | **15** | -10 | | **20** | **25** | -5 | -7 | -3.07 | -2 |
| X | 0.6 | -3 | 0 | 4.4 | -4.64 | -1.14 | -5.51 | 5.66 | 5.92 | -2 | -2 | -4 | -7 | 3 | | | 12 | 9.3 | -2.43 | -1 |
| M | -2 | -2 | -0 | -4 | **16.49** | -2.4 | -5.23 | 11.42 | 1.87 | -1 | -3 | -6 | -8 | -3 | -5 | -9 | | 5.6 | 1.73 | 0.1 |
| F | -1 | -1 | 0.9 | -3 | 2.22 | 0.08 | -1.93 | 4.95 | 1.53 | -1 | -1 | -3 | -3 | -0 | -2 | -2 | 7.4 | | artifact | 10 |
| Y | -0 | 0.8 | 1.2 | -4 | -3.04 | -1.05 | -1.48 | -3.5 | -2.96 | 6.7 | 0.8 | -3 | -4 | -3 | -1 | -2 | -1 | **29** | | 2.7 |
| W | 3.3 | -1 | 0.2 | -2 | -1.75 | 3.19 | -0.18 | -1.96 | -1.57 | -0 | 5.1 | -2 | -1 | 0.8 | -2 | -1 | -0 | -1 | **20.33** | |

| | | | | | | | | | | | | | | | | | | | | |
|---|---|---|---|---|---|---|---|---|---|---|---|---|---|---|---|---|---|---|---|---|
| Chi2 | 89 | 2.8 | 87 | 94 | **215.5** | **297.3** | 2.42 | **436.8** | 48.45 | 0.8 | **427** | 3.2 | 1.5 | 4.8 | 11 | 3.5 | 2.1 | 146 | 56.99 | 16 |

The upper matrix of values are the Z-scores determined by comparing the observed number of substitutions to the distribution of 10,000 simulated numbers of expected substitutions, dependent on marginal probabilities of sources and destinations. The top 5% of Z-scores are bolded. The lower matrix represents the chi-squared values of the translation error rates associated with each amino acid. The top 4 values are bolded.

**A**

Source Amino Acid (columns); Destination Amino Acid (rows)

| | R | H | K | D | E | S | T | N | Q | C | G | P | A | V | I | L | M | F | Y | W |
|---|---|---|---|---|---|---|---|---|---|---|---|---|---|---|---|---|---|---|---|---|
| R | | 3.03 | 6.79 | 0.00 | 0.28 | 7.42 | 0.73 | 0.34 | 0.18 | 0.00 | 13.00 | 0.28 | 0.64 | 0.28 | 2.23 | 1.10 | 0.72 | 0.70 | 6.65 | 0.00 |
| H | 2.38 | | 1.11 | 4.19 | 0.00 | 3.70 | 0.72 | 1.93 | 1.27 | 0.00 | 1.39 | 0.00 | 0.00 | 4.08 | 11.06 | 10.33 | 2.54 | 1.03 | 0.00 | 0.00 |
| K | 0.00 | 14.99 | | 0.97 | 5.18 | 16.79 | 1.21 | 0.53 | 12.55 | 0.00 | 9.05 | 0.93 | 1.31 | 2.62 | 0.28 | 1.14 | 7.11 | 0.00 | 0.00 | 0.00 |
| D | 3.74 | 0.00 | 3.57 | | 14.72 | | 38.19 | | 0.19 | 30.50 | 0.00 | 1.34 | 1.21 | 0.69 | 7.95 | 8.14 | 13.44 | 0.00 | 2.08 | 0.00 |
| E | 0.00 | 1.39 | 7.30 | 1.72 | | 3.92 | | 2.27 | | 5.27 | 0.00 | 11.18 | 2.21 | 3.41 | 0.69 | 0.64 | 12.37 | 22.48 | 5.62 | 14.44 |
| S | 4.24 | 0.00 | 2.46 | 0.00 | | | 20.25 | 0.74 | 6.85 | 9.55 | 1.85 | 7.55 | 39.17 | 0.41 | 0.00 | 0.11 | 1.97 | 2.78 | 0.00 | 6.67 |
| T | 5.89 | 24.42 | 0.00 | 5.33 | 0.13 | 5.68 | | 6.44 | 2.37 | 3.31 | 2.67 | 12.78 | 0.34 | 24.49 | 1.52 | 1.64 | 0.00 | 3.07 | 7.70 | 0.00 |
| N | 4.68 | 1.39 | 1.35 | 30.78 | 15.09 | 12.68 | 1.45 | | 16.77 | 5.56 | 10.22 | 20.57 | 1.01 | 3.65 | 10.59 | 9.08 | 16.10 | 15.97 | 7.68 | 0.00 |
| Q | 2.55 | 2.78 | 33.29 | 1.01 | 16.20 | 9.39 | 2.43 | 4.69 | | 0.00 | 7.19 | 0.48 | 1.38 | 4.38 | 5.57 | 3.98 | 7.35 | 6.55 | 12.88 | 3.33 |
| C | 8.31 | 21.93 | 1.59 | 1.32 | 2.59 | 3.70 | 0.74 | 0.53 | 5.49 | | 2.31 | 1.61 | 0.33 | 5.23 | 11.61 | 17.16 | 1.30 | 0.66 | 38.08 | 0.00 |
| G | 0.61 | 0.00 | 5.75 | 15.45 | 0.26 | 18.77 | 9.92 | 16.21 | 0.74 | 21.64 | | 1.72 | 39.98 | 6.99 | 0.42 | 0.11 | 6.71 | 0.00 | 2.90 | 0.00 |
| P | 18.31 | 10.57 | 0.00 | 2.27 | 1.82 | 3.70 | 4.38 | 5.86 | 0.57 | 0.78 | 2.31 | | 9.07 | 22.45 | 4.32 | 3.49 | 0.00 | 1.06 | 0.00 | 0.00 |
| A | 0.76 | 4.76 | 3.06 | 3.41 | 7.93 | | 3.17 | 8.56 | 8.35 | 2.74 | 24.00 | 3.19 | | 10.34 | 4.86 | 4.17 | 6.21 | 0.00 | 0.00 | 0.00 |
| V | 20.91 | 4.42 | 0.79 | 3.05 | 0.00 | 0.00 | 13.14 | 8.38 | 1.46 | 0.00 | 1.28 | 36.46 | 0.54 | | 35.82 | 37.47 | 0.67 | 0.00 | 0.00 | 0.00 |
| X | 11.43 | 2.78 | 17.54 | 22.70 | 1.34 | 3.51 | 0.98 | 11.57 | 23.15 | 3.92 | 0.00 | 0.76 | 1.55 | 3.99 | | | 10.35 | 0.64 | 0.00 | 0.00 |
| M | 7.51 | 2.78 | 10.44 | 7.02 | 30.73 | 10.73 | 1.71 | 28.98 | 15.43 | 1.55 | 2.67 | 0.96 | 1.17 | 4.22 | 1.55 | 0.43 | | 19.81 | 10.40 | 14.44 |
| F | 1.19 | 0.00 | 1.35 | 0.78 | 2.37 | 0.00 | 0.49 | 2.96 | 3.71 | 5.56 | 0.00 | 0.00 | 0.00 | 1.66 | 0.28 | 0.59 | 11.72 | | | 61.11 |
| Y | 3.74 | 4.76 | 1.67 | 0.00 | 1.06 | 0.00 | 0.49 | 0.00 | 0.56 | 9.64 | 4.06 | 0.00 | 0.10 | 0.97 | 1.11 | 0.22 | 1.45 | 25.24 | | 0.00 |
| W | 3.74 | 0.00 | 1.94 | 0.00 | 0.30 | 0.00 | 0.00 | 0.00 | 0.36 | 0.00 | 17.98 | 0.19 | 0.00 | 0.14 | 0.15 | 0.20 | 0.00 | 0.00 | 6.01 | |

**B**

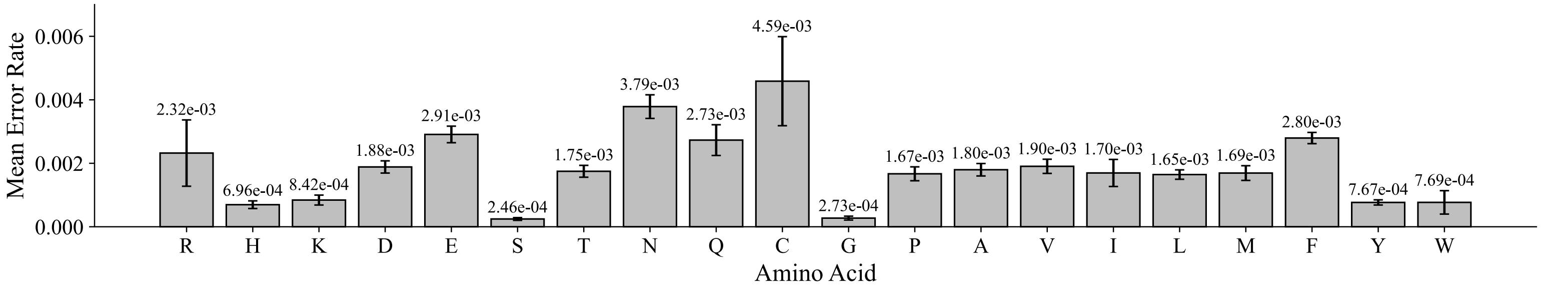


Figure S1. Mean rate and spectrum of translation errors in the accurate ribosomal variant of *E. coli*. A) Wild-type (source) amino acids are represented as columns, and erroneously incorporated (destination) amino acids are represented as rows. B) The mean error rate at which each amino acid is mistranslated is plotted with standard error bars across three biological replicates. Substitutions that are indistinguishable from chemical modifications of the source amino acid (see Table 3) are removed from the analysis (grey boxes). Leucine and isoleucine are combined in the destination row “X.” Subfigure B gives the average rate at which each amino acid is mistranslated, while subfigure A gives the percentage distribution of those mistranslations across destination amino acids.

**A**

Source Amino Acid (columns); Destination Amino Acid (rows)

| | R | H | K | D | E | S | T | N | Q | C | G | P | A | V | I | L | M | F | Y | W |
|---|---|---|---|---|---|---|---|---|---|---|---|---|---|---|---|---|---|---|---|---|
| R | | 0.81 | 9.48 | 0.42 | 0.13 | 5.38 | 0.27 | 0.14 | 0.63 | 0.00 | 5.90 | 0.25 | 0.58 | 0.11 | 1.49 | 0.68 | 0.00 | 0.74 | 6.11 | 0.00 |
| H | 6.90 | | 3.36 | 2.09 | 0.31 | 1.75 | 0.90 | 0.86 | 7.75 | 8.33 | 1.36 | 0.00 | 0.00 | 4.13 | 8.23 | 6.03 | 0.54 | 2.68 | 5.00 | 11.11 |
| K | 1.04 | 6.60 | | 0.95 | 5.82 | 11.26 | 1.89 | 20.38 | 12.00 | 0.00 | 5.78 | 0.43 | 1.86 | 1.80 | 1.38 | 1.01 | 3.01 | 0.00 | 0.00 | 6.06 |
| D | 1.08 | 0.00 | 4.15 | | 17.49 | | 38.76 | | 1.71 | 32.09 | 21.80 | 1.36 | 1.71 | 0.54 | 9.51 | 9.00 | 6.73 | 0.23 | 5.00 | 0.00 |
| E | 0.00 | 0.81 | 10.69 | 38.05 | | 0.00 | | 4.49 | | 5.15 | 5.66 | 10.97 | 0.74 | 3.17 | 0.57 | 1.29 | 9.06 | 15.16 | 2.78 | 3.03 |
| S | 3.92 | 0.00 | 4.15 | 1.52 | | | 17.63 | 0.41 | 9.88 | 11.47 | 7.77 | 7.89 | 40.33 | 0.47 | 0.57 | 0.45 | 1.49 | 1.17 | 0.00 | 11.11 |
| T | 3.16 | 5.66 | 1.96 | 3.89 | 0.00 | 23.40 | | 4.58 | 1.01 | 1.45 | 1.36 | 9.60 | 0.93 | 25.47 | 0.92 | 2.98 | 0.54 | 2.31 | 1.11 | 0.00 |
| N | 3.46 | 3.64 | 6.23 | 19.20 | 11.05 | 17.87 | 2.08 | | 21.24 | 2.78 | 6.02 | 15.97 | 0.69 | 5.14 | 9.12 | 9.02 | 14.34 | 8.78 | 7.78 | 0.00 |
| Q | 6.61 | 55.93 | 27.50 | 1.08 | 15.34 | 3.33 | 0.97 | 3.76 | | 0.00 | 4.43 | 0.85 | 0.41 | 4.22 | 3.56 | 5.36 | 7.96 | 4.99 | 6.39 | 9.70 |
| C | 13.46 | 10.41 | 0.79 | 0.44 | 2.92 | 1.96 | 0.23 | 0.00 | 7.19 | | 0.62 | 2.21 | 0.17 | 3.50 | 12.31 | 14.12 | 3.48 | 0.95 | 27.78 | 0.00 |
| G | 3.17 | 0.00 | 3.97 | 7.85 | 0.34 | 18.98 | 12.65 | 5.46 | 0.47 | 8.05 | | 2.37 | 39.41 | 5.07 | 0.61 | 0.24 | 2.32 | 0.46 | 1.11 | 0.00 |
| P | 24.33 | 6.08 | 1.77 | 1.87 | 2.00 | 0.00 | 4.98 | 5.11 | 0.86 | 0.00 | 0.74 | | 8.71 | 16.95 | 4.15 | 1.87 | 1.39 | 1.20 | 3.61 | 0.00 |
| A | 1.04 | 0.00 | 0.98 | 2.06 | 9.38 | | 3.28 | 9.59 | 5.74 | 5.15 | 26.67 | 4.67 | | 9.96 | 4.88 | 5.08 | 7.08 | 0.00 | 0.00 | 9.70 |
| V | 9.52 | 0.00 | 1.96 | 1.93 | 0.49 | 7.05 | 11.89 | 4.21 | 1.33 | 0.00 | 0.00 | 39.62 | 0.39 | | 36.81 | 40.11 | 1.44 | 0.27 | 0.00 | 0.00 |
| X | 10.62 | 0.00 | 7.10 | 12.15 | 3.14 | 3.92 | 0.45 | 13.64 | 14.05 | 0.00 | 2.94 | 3.39 | 1.65 | 11.31 | | | 30.51 | 20.65 | 0.00 | 0.00 |
| M | 5.28 | 6.20 | 10.69 | 6.28 | 28.28 | 0.00 | 2.09 | 23.22 | 13.25 | 4.23 | 1.60 | 0.43 | 1.86 | 5.82 | 4.41 | 0.47 | | 18.94 | 15.56 | 9.70 |
| F | 1.08 | 0.00 | 1.96 | 0.21 | 2.63 | 1.67 | 0.48 | 4.02 | 2.88 | 0.00 | 0.85 | 0.00 | 0.17 | 1.30 | 0.52 | 1.28 | 8.59 | | | 33.54 |
| Y | 2.15 | 3.85 | 2.75 | 0.00 | 0.54 | 0.00 | 1.01 | 0.14 | 0.00 | 21.30 | 2.71 | 0.00 | 0.00 | 0.28 | 0.94 | 0.62 | 1.08 | 21.24 | | 6.06 |
| W | 3.19 | 0.00 | 0.49 | 0.00 | 0.13 | 3.42 | 0.45 | 0.00 | 0.00 | 0.00 | 3.80 | 0.00 | 0.39 | 0.77 | 0.00 | 0.39 | 0.44 | 0.23 | 17.78 | |

**B**

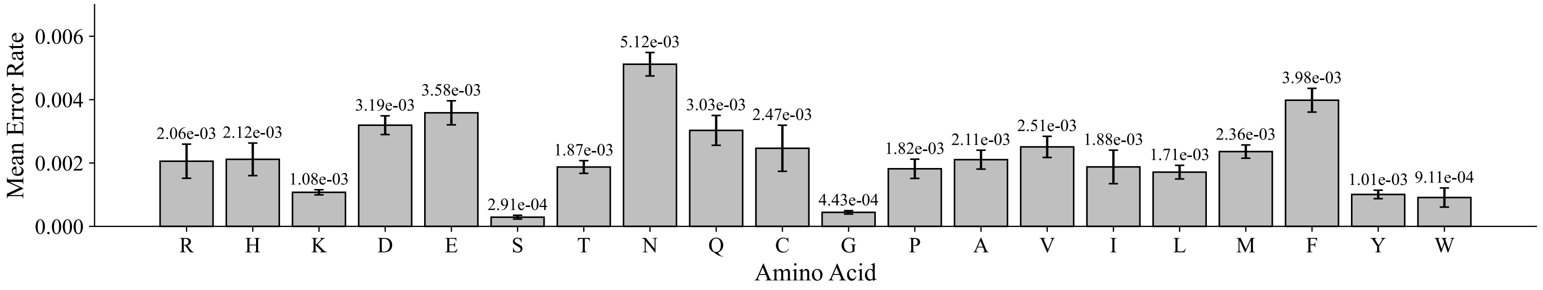


Figure S2. Mean rate and spectrum of translation errors in the error-prone ribosomal variant of *E. coli*. A) Wild-type (source) amino acids are represented as columns, and erroneously incorporated (destination) amino acids are represented as rows. B) The mean error rate at which each amino acid is mistranslated is plotted with standard error bars across three biological replicates. Substitutions that are indistinguishable from chemical modifications of the source amino acid (see Table 3) are removed from the analysis (grey boxes). Leucine and isoleucine are combined in the destination row "X." Subfigure B gives the average rate at which each amino acid is mistranslated, while subfigure A gives the percentage distribution of those mistranslations across destination amino acids.

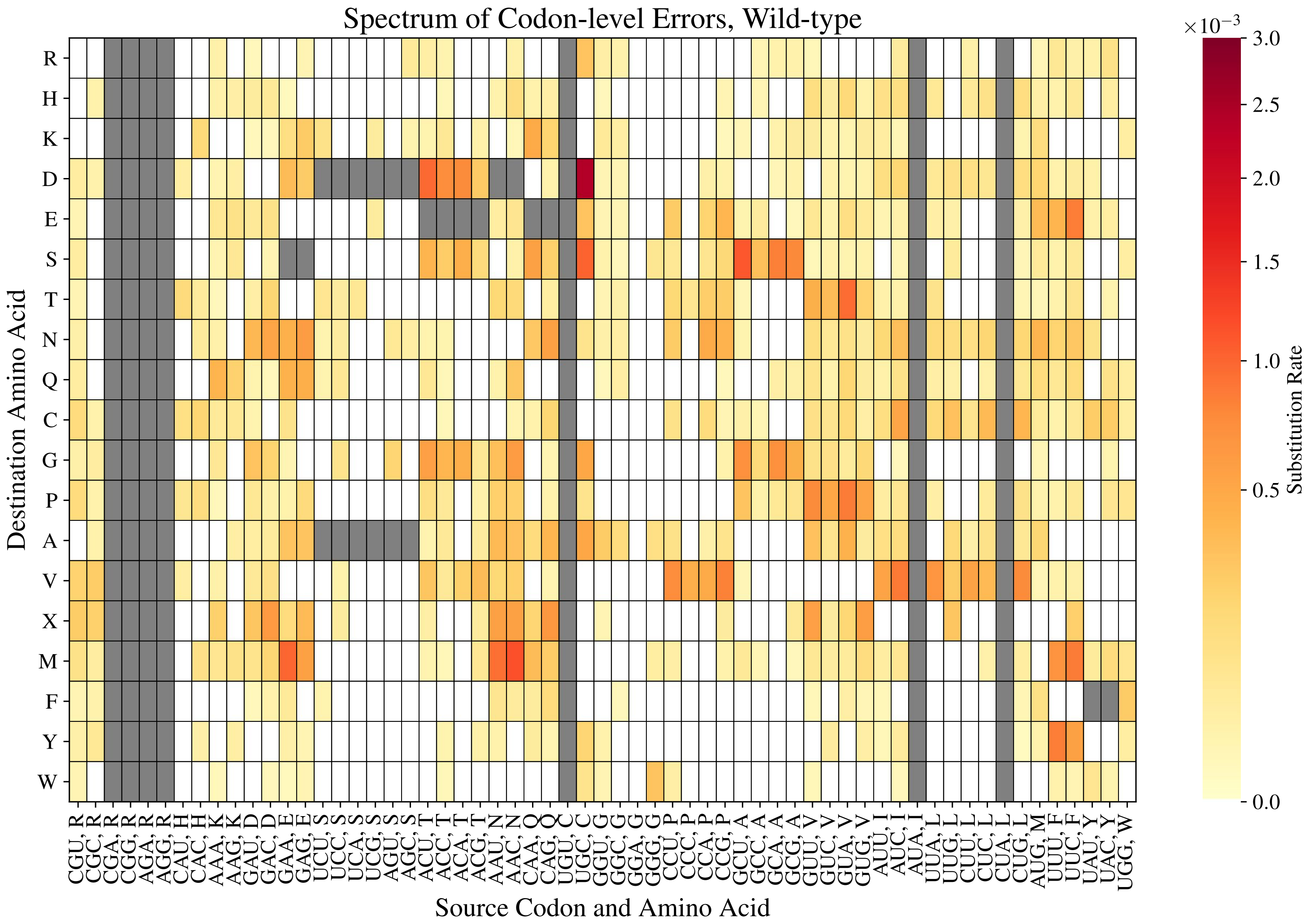


Figure S3. Mean codon-specific substitution rate of the wild-type ribosomal variant of *E. coli*. Codon results are filtered to include only those with at least 3,000 sampled codons. Grey boxes denote that the result was filtered due to either chemical artifact (see Table 3), or failure to meet the sampling cutoff. Substitution bias, used in the calculation for substitution rate, is the fraction of observed errors from a source codon that are directed to a specific destination amino acid. The codon-specific substitution rate is calculated as the product of the mean codon error rate and the mean substitution bias of a source codon toward each possible destination amino acid.

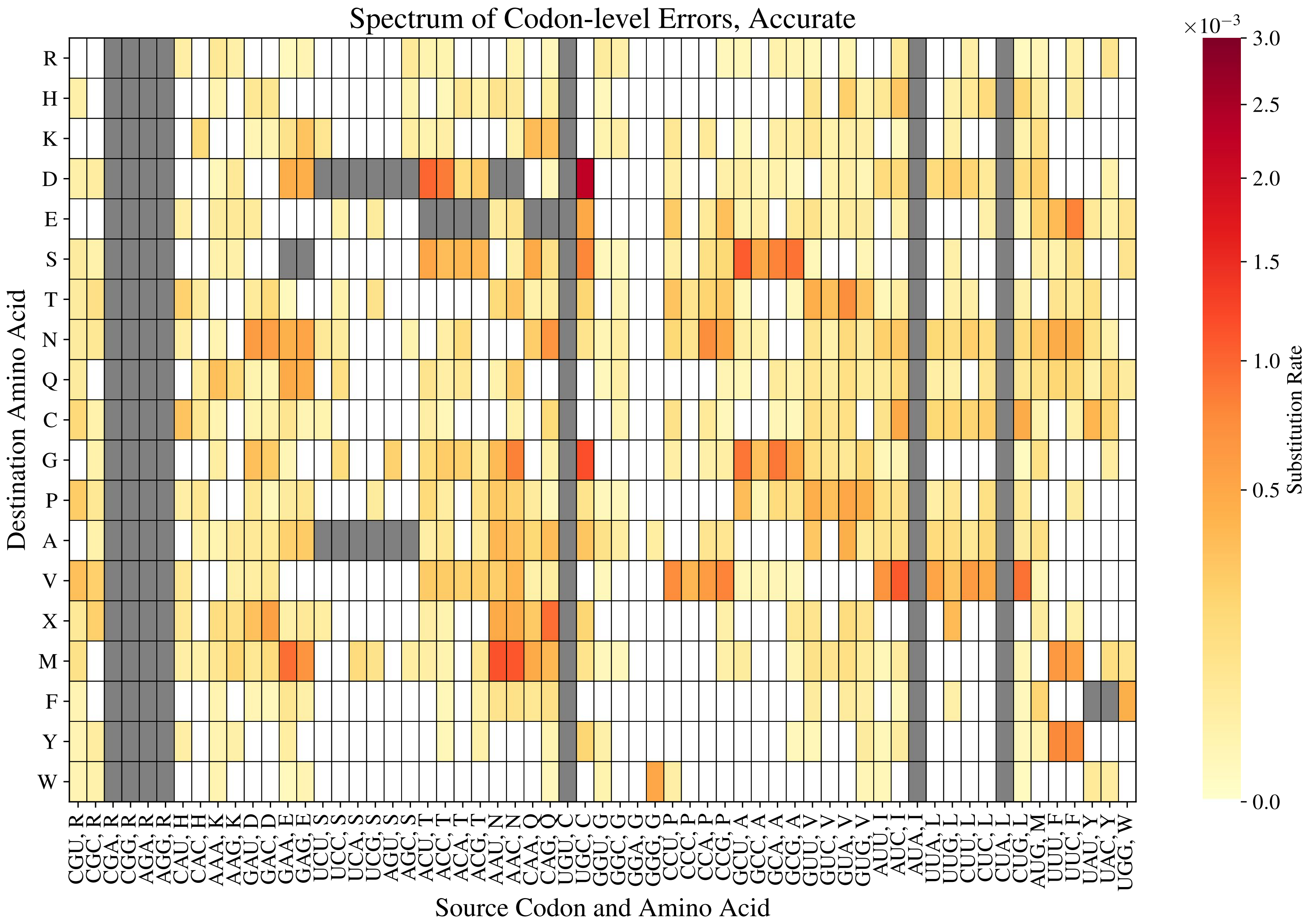


Figure S4. Mean codon-specific substitution rate of the accurate ribosomal variant of *E. coli*. Codon results are filtered to include only those with at least 3,000 sampled codons. Grey boxes denote that the result was filtered due to either chemical artifact (see Table 3), or failure to meet the sampling cutoff. Substitution bias, used in the calculation for substitution rate, is the fraction of observed errors from a source codon that are directed to a specific destination amino acid. The codon-specific substitution rate is calculated as the product of the mean codon error rate and the mean substitution bias of a source codon toward each possible destination amino acid.

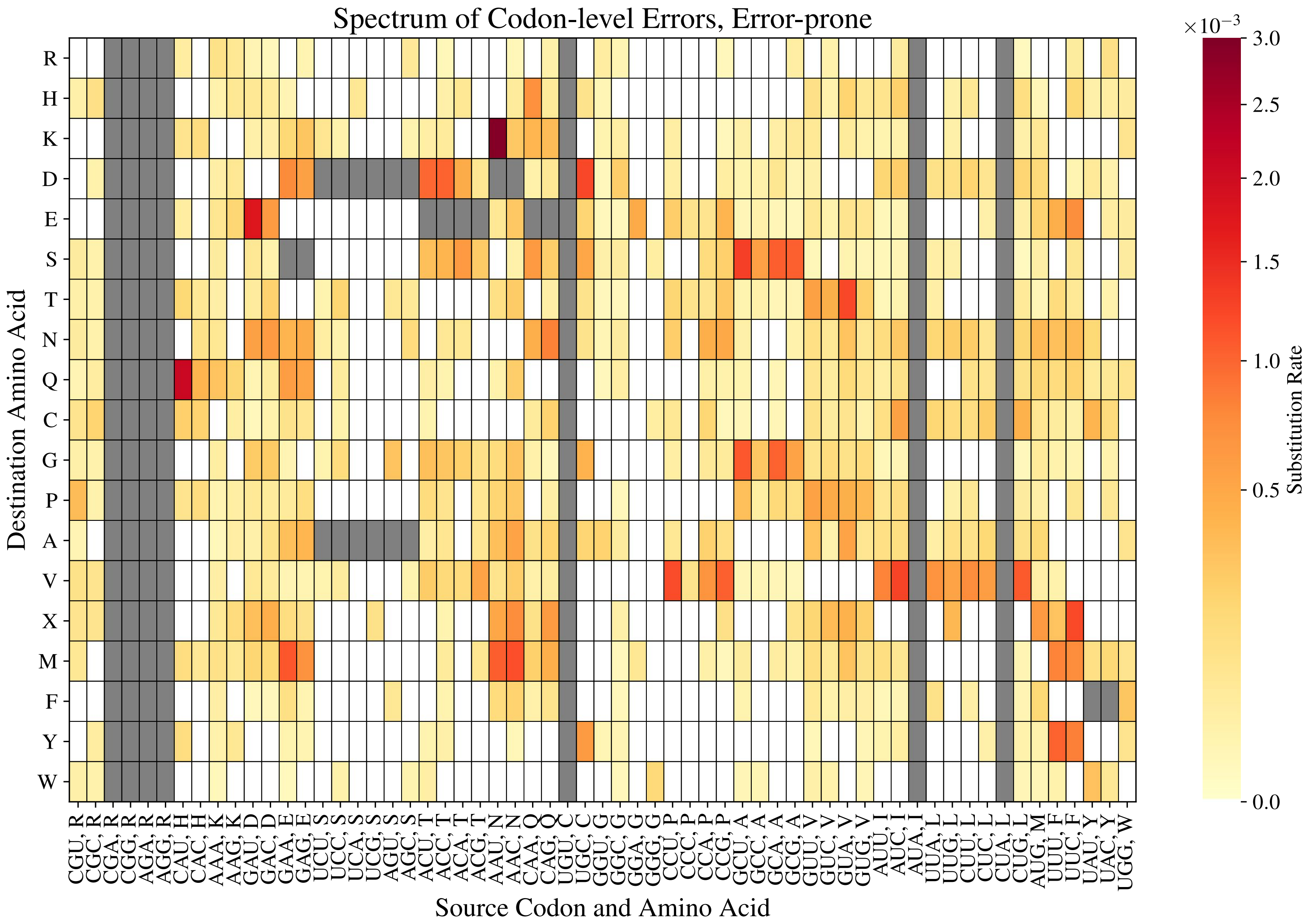


Figure S5. Mean codon-specific substitution rate of the error-prone ribosomal variant of *E. coli*. Codon results are filtered to include only those with at least 3,000 sampled codons. Grey boxes denote that the result was filtered due to either chemical artifact (see Table 3), or failure to meet the sampling cutoff. Substitution bias, used in the calculation for substitution rate, is the fraction of observed errors from a source codon that are directed to a specific destination amino acid. The codon-specific substitution rate is calculated as the product of the mean codon error rate and the mean substitution bias of a source codon toward each possible destination amino acid.

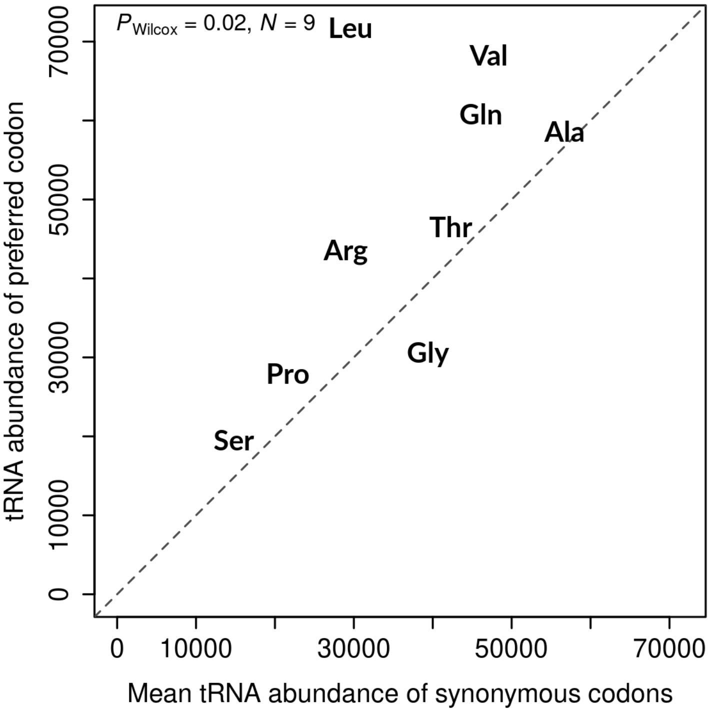


Figure S6. Preferred codons correspond to more abundant tRNA species. The tRNA abundance of the preferred codon for each amino acid is greater than the mean tRNA abundance over its synonymous codons for all amino acids except Gly. This analysis was done for nine amino acids with available tRNA abundance data from [1] for more than one iso-accepting tRNA species, which were identified on the basis of the revised wobble rules for the four bases – A,C,G, and U[2].

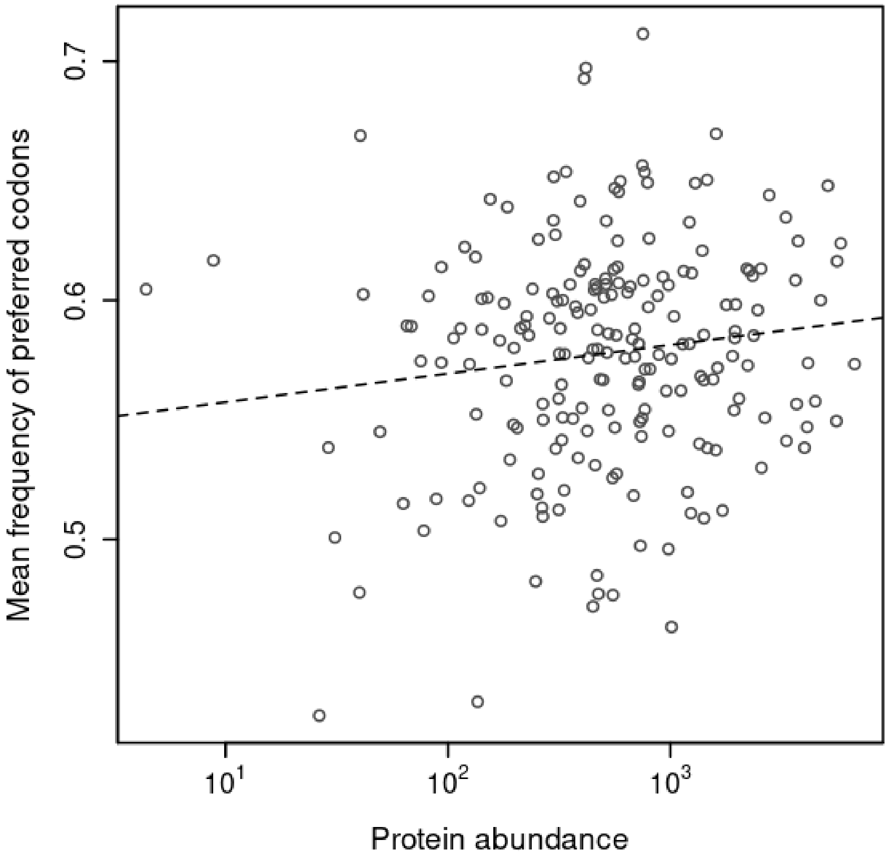


Figure S7. The mean frequency of preferred codons among synonymous codons in a gene are uncorrelated with its protein abundance, for the set of preferred codons predicted based on codon frequencies of all genes. The linear regression coefficient for a correlation between this measure of codon usage bias and log protein abundance for 205 genes (data from PaxDb[3]) was $\beta = 0.012$ (SE = 0.006) and the Spearman rank correlation coefficient was $\rho = 0.102$ ($P = 0.15$). This figure is comparable to main Figure 3A which shows an enrichment of preferred codons with increasing protein abundance, for the set of preferred codons predicted based on codon frequencies of highly expressed genes.

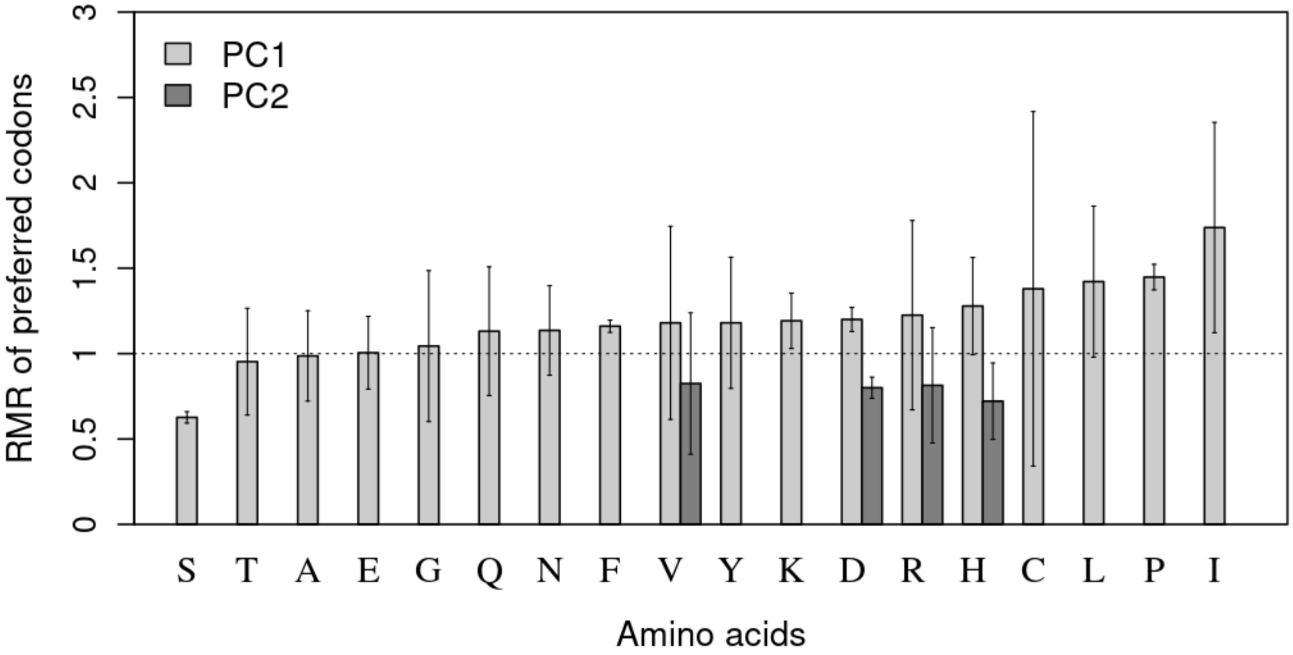


Figure S8. RMR of preferred codons of 18 amino acids, for the two sets of preferred codons - predicted based on codon frequencies of highly expressed genes (PC1), or of all genes (PC2). Preferred codons, that differ between these two sets for four amino acids, have lower mean RMR in PC2. The error bars denote one standard error of the mean RMR over three biological replicates.

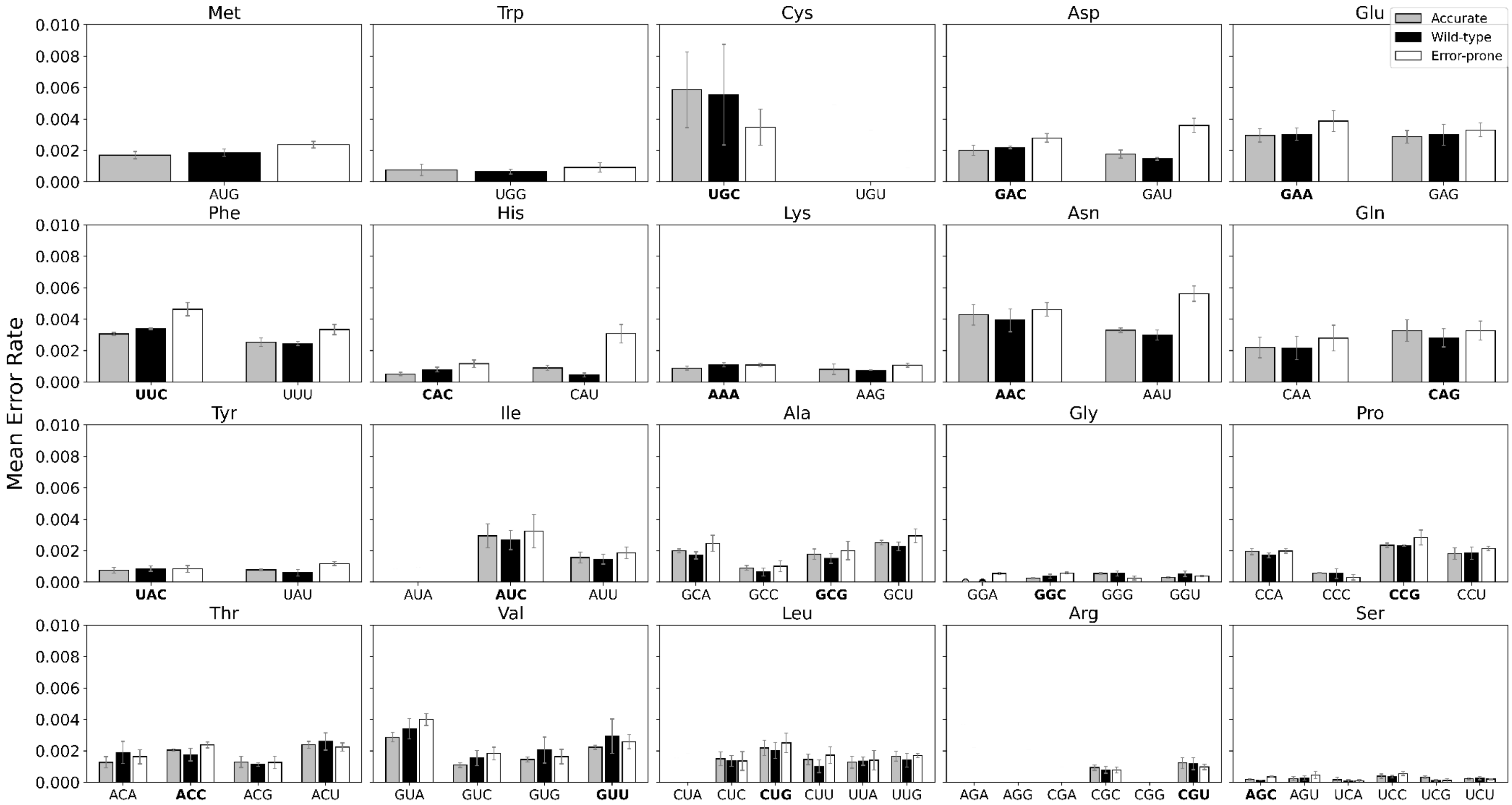


Figure S9. Mean error rates of each codon according to the ribosomal variant of *E. coli*. A minimum of 3,000 sampled codons is required per each plotted result. Mean error rates are plotted here with standard error bars across three biological replicates. Codons with measured error rates of zero are plotted as points (e.g. GGA). Preferred codons are indicated in bold. Ribosomal variants are distinguished by the colors shown in the legend.

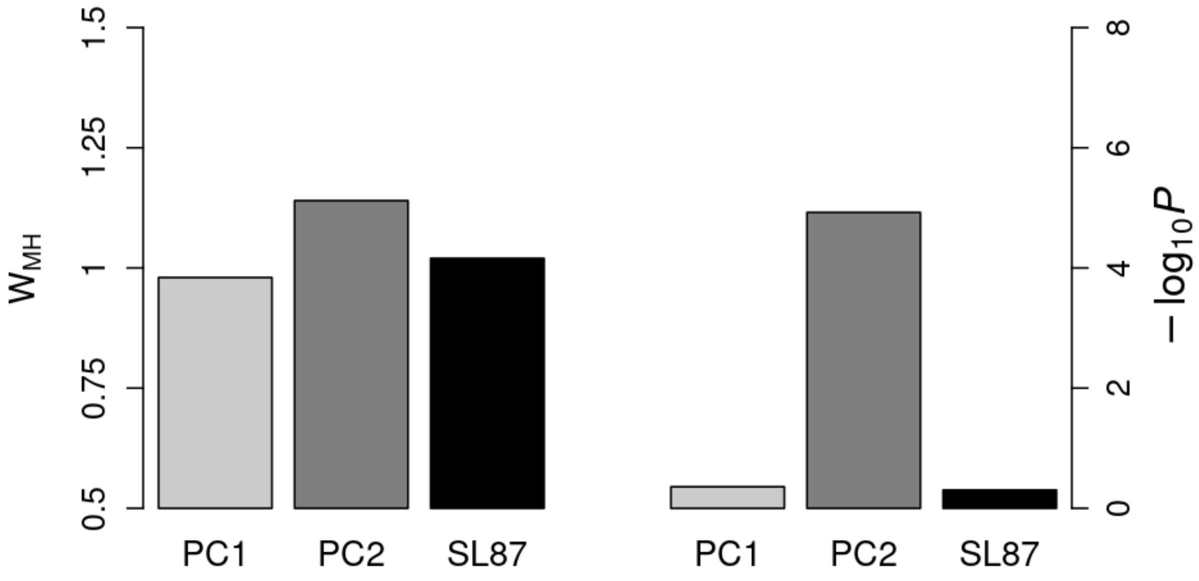


Figure S10. Akashi's test of enrichment of preferred codons at conserved amino-acid sites[4] for three sets of preferred codons. PC1 and PC2 are predicted based on codon frequencies of highly expressed genes and of all genes, respectively. SL87 is the set of preferred codons predicted by Sharp and Li (1987)[5]. WMH is the Mantel-Haenszel's statistic for the enrichment of preferred codons at conserved sites relative to variable sites (Methods)[6]. This is equivalent to an Odds Ratio (OR) for the counts of preferred and non-preferred synonymous codons pooled across different amino acids and genes. WMH > 1 indicates an enrichment of preferred codons at conserved sites. Log-transformed p-values are shown to the right, where a value greater than 2 indicates a significant result ($P < 0.01$).
